# NHR-23 activity is necessary for *C. elegans* developmental progression and apical extracellular matrix structure and function

**DOI:** 10.1101/2021.10.27.465992

**Authors:** Londen C. Johnson, An A. Vo, John C. Clancy, Krista M. Myles, Murugesan Pooranachithra, Joseph Aguilera, Max T. Levenson, Chloe Wohlenberg, Andreas Rechtsteiner, James Matthew Ragle, Andrew D. Chisholm, Jordan D. Ward

**Affiliations:** Department of Molecular, Cell, and Developmental Biology, University of California-Santa Cruz, Santa Cruz, CA 95064, USA.; Department of Cell and Developmental Biology, School of Biological Sciences, University of California San Diego, La Jolla, CA 92093, USA.

**Keywords:** *C. elegans*, molting, NHR-23, nuclear hormone receptor, apical extracellular matrix, auxin-inducible degron

## Abstract

This work shows how a *C. elegans* transcription factor controls remodeling of the apical extracellular matrix during development and in which tissues it acts.

**ABSTRACT:** Nematode molting is a remarkable process where animals must repeatedly build a new apical extracellular matrix (aECM) beneath a previously built aECM that is subsequently shed. The nuclear hormone receptor NHR-23/NR1F1 is an important regulator of *C. elegans* molting. NHR-23 expression oscillates in the epidermal epithelium, and soma-specific NHR-23 depletion causes severe developmental delay and death. Tissue-specific RNAi suggests that *nhr-23* acts primarily in seam and hypodermal cells. NHR-23 coordinates the expression of factors involved in molting, lipid transport/metabolism, and remodeling of the aECM. NHR-23 depletion causes dampened expression of a *nas-37* promoter reporter and a loss of reporter oscillation. The cuticle collagen ROL-6 and zona pellucida protein NOAH-1 display aberrant annular localization and severe disorganization over the seam cells following NHR-23 depletion, while the expression of the adult-specific cuticle collagen BLI-1 is diminished and frequently found in patches. Consistent with these localization defects, the cuticle barrier is severely compromised when NHR-23 is depleted. Together, this work provides insight into how NHR-23 acts in the seam and hypodermal cells to coordinate aECM regeneration during development.

## INTRODUCTION

Molting is a critical developmental process required for the growth of all ecdysozoans, a clade comprising an estimated total of 4.5 million living species (Telford et al., 2008). Invertebrate molting involves conserved processes such as apical extracellular matrix (aECM) remodeling, intracellular trafficking, and oscillatory gene expression (Lažetić and Fay, 2017). Molting is also a process of interest for developing new drugs against parasitic nematodes (Ghedin et al., 2007). Parasitic nematodes cause over a billion human infections each year and loss of livestock and crops (Ward, 2015b). However, very few drugs exist, and resistance to those drugs is emerging rapidly (Ward, 2015b). While many genes involved in nematode molting have been identified in *C. elegans* (Frand et al., 2005), little is known about how their gene products are coordinated to promote aECM remodeling and the generation and release of a new cuticle.

Nematodes progress through four periodic larval stages (L1-L4), prior to becoming a reproductive adult (Brenner, 1974). The end of each larval stage is punctuated by a molt that involves trafficking and secretion of aECM components, assembly of a new aECM underneath the old cuticle, followed by separation of the cuticle from the underlying aECM (apolysis) and shedding of the old cuticle (ecdysis) (Lažetić and Fay, 2017). Apolysis coincides with a sleep-like behavior called lethargus (Singh and Sulston, 1978). The cuticle is a collagenous exoskeleton secreted by hypodermal and seam epithelial cells (Page and Johnstone, 2007). The outermost layer (glycocalyx) is rich in carbohydrates and mucins (Nelson et al., 1983; Singh and Sulston, 1978). Beneath the glycocalyx is the glycolipid- and lipid-rich epicuticle which is postulated to function as a hydrophobic surface barrier (Blaxter, 1993; Blaxter et al., 1992). Underlying the epicuticle is a layer comprised mainly of cuticlin proteins. Between the epidermal membrane and the epicuticle are collagen rich layers, including a fluid-filled medial layer with collagen struts (Lažetić and Fay, 2017; Page and Johnstone, 2007). Sets of collagens oscillate in expression over the course of each molt and are classified into three groups based on temporal expression: early, intermediate, and late collagens (Johnstone and Barry, 1996; Page and Johnstone, 2007). While the specific localization of most collagens is unknown, collagens from the same group are thought to be in the same layer (Lažetić and Fay). A transient structure, the sheath or pre-cuticle, is formed during each larval stage and thought to pattern the cuticle (Cohen and Sundaram, 2020). Many of the components of this pre-cuticle are related to mammalian matrix proteins. The *C. elegans* pre-cuticle contains Zona Pellucida proteins, proteins related to small leucine-rich proteoglycans, and lipid transporters in the lipocalin family (Cohen and Sundaram, 2020; Cohen et al., 2020; Forman-Rubinsky et al., 2017; Kelley et al., 2015; Sapio et al., 2005; Vuong-Brender et al., 2017).

Nuclear hormone receptor (NHR) transcription factors are key regulators of molting in insects and nematodes (King-Jones and Thummel, 2005; Taubert et al., 2011). NHRs are characterized by a ligand-binding domain (LBD) which has the potential to bind small molecules such as ligands and dietary-derived metabolites (Taubert et al., 2011). NHRs have a canonical zinc finger DNA-binding domain (DBD) with an unstructured hinge region between the DBD and LBD that is subject to post-translational regulation (Antebi, 2015; Campbell et al., 2008; Ward et al., 2013). A single, conserved nuclear hormone receptor, NHR-23/NR1F1 (hereafter referred to as NHR-23), an ortholog of DHR3 in insects and ROR in mammals, is a key regulator of *C. elegans* molting. NHR-23 is also necessary for spermatogenesis (Ragle et al., 2020; Ragle et al., 2022). *nhr-23* mutation or inactivation by RNAi leads to embryonic lethality, larval arrest, ecdysis defects, and morphology defects (Frand et al., 2005; Gissendanner et al., 2004; Kostrouchova et al., 1998; Kostrouchova et al., 2001). *nhr-23* mRNA expression oscillates over the course of each larval stage, peaking mid-larval stage and hitting a trough at the molt (Gissendanner et al., 2004; Kostrouchova et al., 2001). *nhr-23* is necessary for all four larval molts and it regulates microRNAs such as *let-7* and *lin-4* (Kostrouchova et al., 1998; Kostrouchova et al., 2001; Patel et al., 2022, Kinney et al., 2023). *let-7* also regulates *nhr-23,* suggesting that a feedback loop might coordinate molting with developmental timing (Patel et al., 2022). The NHR-23 insect ortholog (DHR3) is part of the molting gene regulatory network (Lam et al., 1997; Ruaud et al., 2010) and the mammalian ortholog (RORα) regulates circadian rhythms, lipid metabolism, and immunity (Jetten, 2009). However, how NHR-23 regulates molting and whether it is part of the core oscillator that promotes rhythmic gene expression over each larval stage is poorly understood (Tsiairis and Großhans, 2021).

We show here that NHR-23 protein oscillates and rapid NHR-23 depletion via an auxin-inducible degron causes severe developmental delay and death. Analysis of NHR-23 target genes suggests a role in coordinating aECM assembly, remodeling, and construction of specific cuticular structures. NHR-23 depletion causes aberrant localization of the collagens ROL-6 and BLI-1 and of the pre-cuticle factor NOAH-1. NHR-23 activity is necessary in seam and hypodermal cells to promote molting and timely development. Our work reveals when and where NHR-23 acts to promote molting.

## RESULTS

### NHR-23 protein oscillates during development

Expression of *nhr-23* mRNA oscillates throughout each *C. elegans* larval stage (Gissendanner et al., 2004; Hendriks et al., 2014; Kostrouchova et al., 2001; Meeuse et al., 2020). However, mRNA expression profiles do not always correlate with protein levels (de Sousa Abreu et al., 2009; Vogel and Marcotte, 2012). To determine whether NHR-23 protein oscillates, we monitored the expression of an endogenously tagged NHR-23::GFP, over 28 hours (*wrd8* allele; Ragle et al., 2020). During L1, oscillating expression of NHR-23::GFP was observed in the nuclei of seam and hypodermal cells (Fig. S1). Expression of NHR-23 rises and falls prior to the completion of the 1^st^ molt (Fig. 1A). As GFP tags can affect the expression or stability of proteins (Agbulut et al., 2006; Baens et al., 2006), we assayed the expression of NHR-23 through western blotting time-courses using an *nhr-23::AID*::3xFLAG* strain (*kry61* allele; Zhang et al., 2015). These experiments confirmed that NHR-23::AID*::3xFLAG also oscillates (Fig. 1B, C), similar to our NHR-23::GFP imaging experiments (Fig. 1A). We detected three distinct NHR-23 bands; the lowest band is consistent with the size of NHR-23b/f (Fig. 1B, C, Fig. S2). The upper two bands are larger than expected, which could reflect post-translational modification or that NHR-23 migrates aberrantly during SDS-PAGE. All observed NHR-23 bands oscillate, though the lowest band was expressed more strongly than the other bands (Fig. 1B, C). The peak in NHR-23 expression is earlier in each larval stage than previous qRT-PCR approaches suggested (Gissendanner et al., 2004; Kostrouchova et al., 2001) and more in line with recent RNA-seq and imaging data (Meeuse et al., 2020; Kinney et al., 2023). To understand how NHR-23 promotes molting, it is important to clarify when it peaks in expression. We examined NHR-23::GFP expression over the course of the 4^th^ larval stage, as vulva morphology allows for more precise sub-staging (Mok et al., 2015). NHR-23::GFP expression was first detectable in L4.1 larvae, peaked in expression from L4.2-L4.3, and disappeared in the vulva by L4.5 and head by L4.6 and then remained undetectable (Fig. 1D,E, S1B,C), similar to what Kinney et al. (2023) observed with NHR-23::mScarlet.

**Fig. 1.**
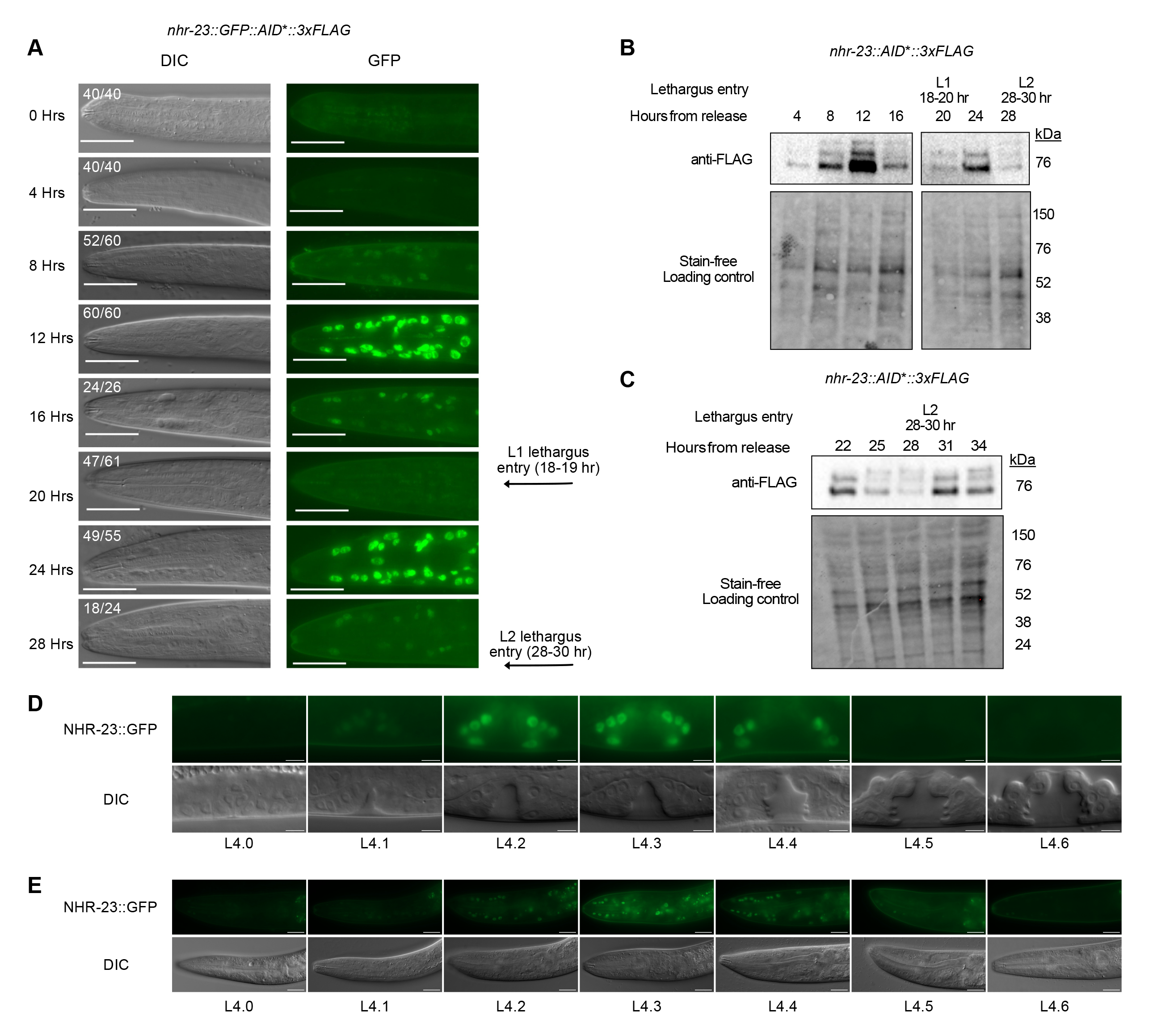
NHR-23 protein oscillates during development. (A) Representative images from a timecourse monitoring the expression of endogenous NHR-23::GFP protein Two biological replicates were performed and the number of animals for which the image is representative at each timepoint is provided. Scale bars=20 µm. (B and C) Anti-FLAG immunoblot analysis of synchronized *nhr-23::AID*\*::3xFLAG animals monitoring the expression of NHR-23 3XFLAG across two timecourses, 4-28 hours (B) and 22-34 hours (C). Marker size (in kDa) is provided. Stain-free analysis, which visualizes total protein on the membrane, is provided as a loading control. The blots are representative of three experimental replicates. The times at which animals were observed to enter lethargus is indicated in A-C. NHR-23::GFP protein expression in the vulva (D) and head (E) during the L4 larval stage. Animals were staged based on vulval morphology (Mok et al., 2015). Images are representative of 20 animals examined over 4 independent experiments. Scale bars=5 µm (D) or 20 µm (E). *nhr-23::GFP::AID*::3xFLAG* (A,C,E) and *nhr-23::AID*::3xFLAG* (B,C) are previously described endogenous knock-ins that produce C-terminal translational fusions to all known *nhr-23* isoforms (Ragle et al., 2020; Zhang et al., 2015).

### NHR-23 depletion causes severe developmental delay

Given that NHR-23 oscillates, we tested when NHR-23 was necessary for molting during the L1 larval stage. In a pilot experiment, we consistently found weaker depletion phenotypes using an *nhr-23::GFP::AID*::3xFLAG* strain (Ragle et al., 2020) compared to the *nhr-23::AID*::3xFLAG strain* (Zhang et al., 2015) (Table S1). We therefore transitioned to using the *nhr-23::AID*::3xFLAG* strain for all experiments involving phenotypic analysis. We first tested NHR-23::AID*::3xFLAG depletion kinetics and found robust depletion within 15 minutes after exposure to 4 mM auxin; levels remained low over the 16 hour timecourse (Fig. 2A).

**Fig. 2.**
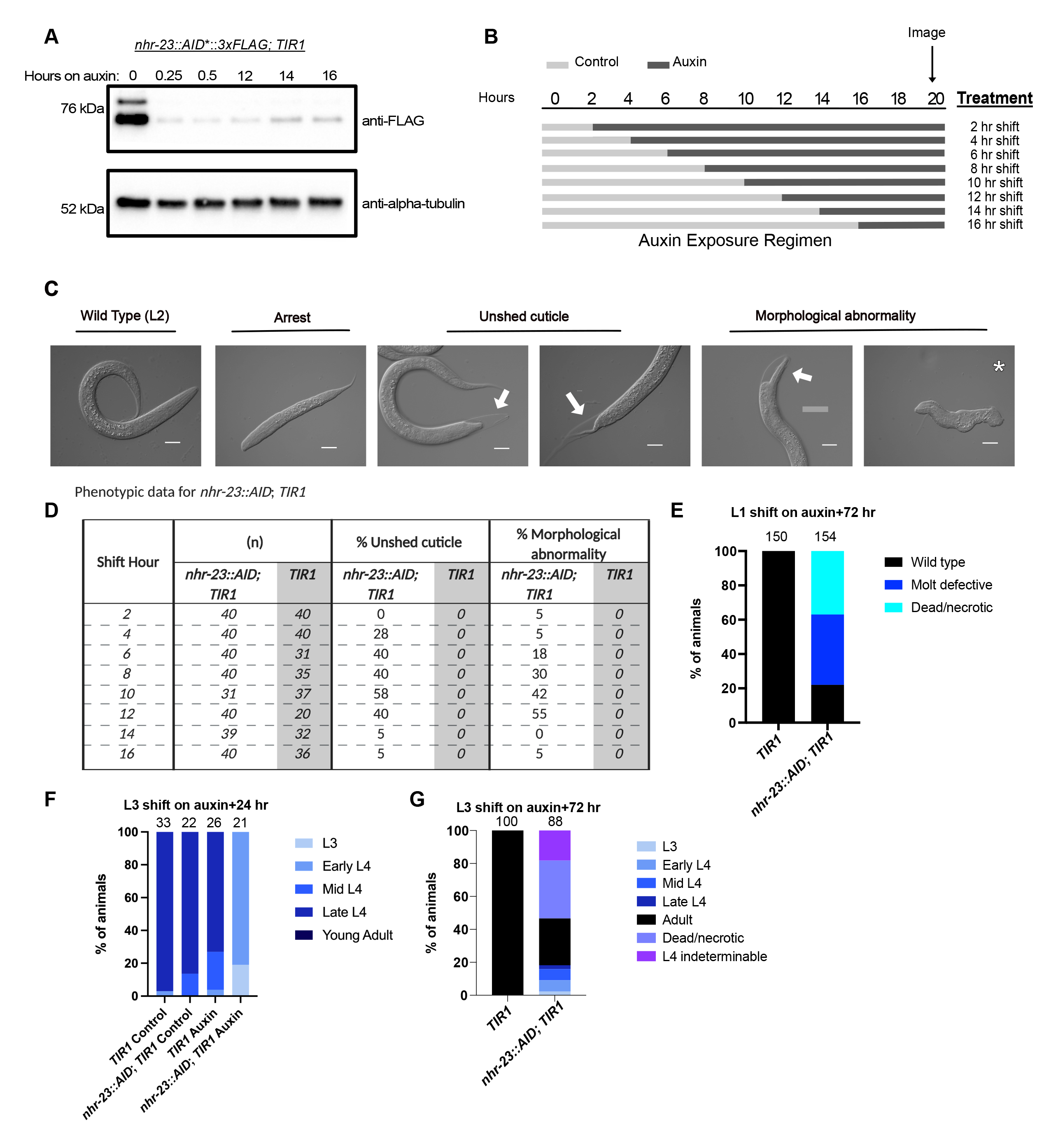
Distinct phenotypes are observed when NHR-23 is depleted early or late in L1 larvae. (A) Anti-FLAG immunoblot to monitor NHR-23 depletion. Synchronized *nhr-23::AID*::3xFLAG; TIR1* and *TIR1* animals were grown on MYOB plates for 12 hours and then shifted onto control or auxin plates. Lysates were collected at the indicated time points. Marker size (in kDa) is provided. An anti-alpha-tubulin immunoblot is provided as a loading control. The blots are representative of three experimental replicates. (B) Schematic of the experimental set-up. Synchronized L1 larvae were grown on control media (light grey bands) and transferred to 4 mM auxin (dark grey bands) every 2 hours. Animals remained on auxin until they were imaged at 20 hours post-release. (C) Representative images of phenotypes observed when *nhr-23::AID*::3xFLAG; TIR1* animals were exposed to auxin in the first larval stage. (D) Percent of animals of the indicated genotypes with unshed cuticles and morphological abnormalities following exposure to auxin. The number of animals scored (n) is provided. (E) Synchronized *nhr-23::AID; TIR1* L1 larvae were released on auxin and scored for molting defects and death 72 hours post-release. (F) Animals of the indicated genotype were synchronized and shifted onto auxin at 25 hours post-release and imaged 23 hours later (48 hours post-release). Animals were staged based on vulval morphology (Mok et al., 2015). L4 larval stages were grouped as Early L4 (4.0-4.2), Mid L4 (4.3-4.5), and Late L4 (4.6-4.9). (G) *nhr-23::AID; TIR1* animals were treated as in (F) and scored after 72 hours on auxin. The number of animals assayed for each genotype and condition is indicated at the top of the graph in E-G.

We performed timed depletion experiments in the L1 larval stage (Fig. 2B) using an *eft-3p::TIR1::mRuby2* control strain (henceforth referred to as *TIR1*) and a strain that permits somatic depletion of NHR-23 in the presence of auxin, (*eft-3p::TIR1::mRuby2; nhr-23::AID*::3xFLAG;* henceforth referred to as *nhr-23::AID*; *TIR1*). L1s were synchronized by starvation arrest and release onto MYOB plates. Animals were shifted onto either 4 mM auxin or control plates every two hours and scored for molting defects at 20 hours post-release from starvation (Fig. 2B). All *TIR1* control animals shifted onto control or auxin and *nhr-23::AID*; *TIR1* animals shifted onto control plates reached the L2 stage (Fig. 2C, D). In contrast, *nhr-23::AID*; *TIR1* animals shifted onto auxin displayed multiple phenotypes including molting defects, internal vacuoles, developmental abnormalities such as disrupted tails and viable animals with a squashed morphology (Fig. 2C). We then quantified the molting defects, scoring animals unable to shed their cuticles, or with morphological abnormalities (Fig. 2D). Animals shifted onto auxin within the first four hours post-release tended to arrest as L1 larvae with few molting or morphological defects (Fig. 2D). To determine at what stage these animals arrested we used an *hlh-8p::GFP* promoter reporter to monitor the M cell lineage (Harfe et al., 1998). Newly hatched L1 animals have one M cell which undergoes a stereotypical series of divisions to produce 16 M lineage cells by the L1 molt (Fig. S3A) (Sulston and Horvitz, 1977). *nhr-23::AID*; *TIR1* animals shifted onto auxin at 3 hours post-release were late in L1 based on M cell number (Fig. S3B, C). When NHR-23 animals were scored 72 hours post-auxin shift we observed molting defects and necrotic animals (Fig. 2E, S4A), suggestive of a severe developmental delay. We also observed arrested animals with a wild-type morphology which were likely L1 larvae by size (Fig. 2E). Shifts between 6- and 12-hours post-release resulted in increased morphological and molting defects (Fig 2D). Animals shifted onto auxin at 9 hours reached the end of the L1 stage by M cell number (S3B, C). Shifts at 14- and 16-hours post-release resulted in wild-type L2 animals as judged by size (Fig. 2D).

To test whether NHR-23 was similarly required in other larval stages, we performed depletion experiments later in development where we could use vulval morphology to score progression through the L4 stage. We performed depletion experiments shifting early L3 animals onto auxin and monitored development by scoring vulva morphology 23 hours later. The majority of *TIR1* animals grown on control or auxin plates and *nhr-23::AID*; *TIR1* animals grown on control plates reached late L4, with a fraction of the population in early or mid-L4 (Fig. 2F, S4B). In contrast, when *nhr-23::AID*; *TIR1* animals were shifted to auxin, the vast majority of animals were early L4 larvae with a fraction of the population remaining in L3 (Fig. 2F, S4B). Repeating these experiments scoring later timepoints revealed that NHR-23-depleted animals were slowly progressing through development. After 72 hours on auxin we observed adult animals as evidenced by the presence of oocytes (Fig. 2G, S4C) while all control animals were adults. A subset of NHR-23-depleted L4 larvae could not be precisely staged due to aberrant vulval morphology (Fig. S4C). These animals had large vulval lumens reminiscent of a 4.3 stage vulva (Mok et al., 2015), but we also observed adults with a similar large vulval lumen (Fig. S4C). A fraction of *nhr-23::AID*; *TIR1* animals grown for 72 hours on auxin were dead and appeared necrotic with large fluid filled spaces within the animal (Fig. 2G, S4C). These depletion experiments indicate that NHR-23 is required for timely developmental progression through multiple larval stages, consistent with previous reports of *nhr-23* inactivation by RNAi (Macneil et al., 2013; Patel et al., 2022)

### NHR-23 regulates oscillating genes involved in aECM biogenesis and regulation of protease activity

To gain insight into how NHR-23 depletion causes developmental delay, we analyzed the expression of *nhr*-23-regulated genes by mining an existing microarray (Kouns et al., 2011) and oscillatory gene expression datasets (Meeuse et al., 2020) (Table S2). Of the 265 *nhr-23-*regulated genes in the microarray dataset, 236 (89%) were oscillatory with the bulk of genes peaking in expression between 180° and 360° (Fig. 3A, Table S2); *nhr-*23 mRNA peaks at 178.11° (Meeuse et al., 2020). A similar trend was observed in a recent analysis of these datasets (Tsiairis and Großhans, 2021). In contrast, 10-20% of *C. elegans* genes oscillate in expression (Hendriks et al., 2014; Kim et al., 2013; Meeuse et al., 2020) (Two-tailed Chi-squared test p<0.0001). *nhr-23-* regulated oscillating genes were enriched in gene ontology classes such as cuticle structure, regulation of endopeptidases, and metalloprotease activity (Table S3). These gene ontology classes are a specific subset of the functions enriched in all oscillating genes (Table S4). To provide biological context, we converted the 360° period to developmental time assuming a 9-hour larval stage and set the molt from the start of lethargus (45°) to the end of ecdysis (135°) as in Meeuse et al. (2023). Most *nhr-23*- regulated genes involved in aECM structure/function, cholesterol metabolism, molting regulation, transcriptional regulation, and signal transduction had peak amplitudes within three hours of the *nhr-23* expression peak (Fig. 3B). Genes involved in the blocked protein unfolding response, as well as some aECM genes peaked later in each larval stage (Fig. 3B). While the *C. elegans* genome encodes 181 collagen genes (Teuscher et al., 2019), *nhr-23* only regulated 13 collagens including all of the furrow collagens implicated in epidermal damage sensing; these furrow collagens are in the “early” collagen group (*dpy-2, dpy-3, dpy-7, dpy-10;* Table S2, Fig. 3C) (Johnstone and Barry, 1996; Page and Johnstone, 2007; Dodd et al., 2018). Other *nhr-23-*regulated intermediate collagens have well-described roles in body morphology (*sqt-1, sqt-2, rol-6*; Table S2, Fig. 3C) (Kramer and Johnson, 1993; Johnstone and Barry, 1996; Page and Johnstone, 2007). There are also several sets of genes involved in aECM biogenesis and remodeling, such as lipocalins, proteases, protease inhibitors, fibrillin, PAN and zona pellucida (ZP) domain-containing proteins, and leucine-rich repeat proteins (Fig. 3C). Although not enriched as a gene ontology class, NHR-23 also regulates several transcription factors implicated in molting or energy metabolism (*peb-1, nhr-91, dpy-20*; Table S2, Fig. 3B) (Clark et al., 1995, 20; Fernandez et al., 2004; Kasuga et al., 2013).

**Fig. 3.**
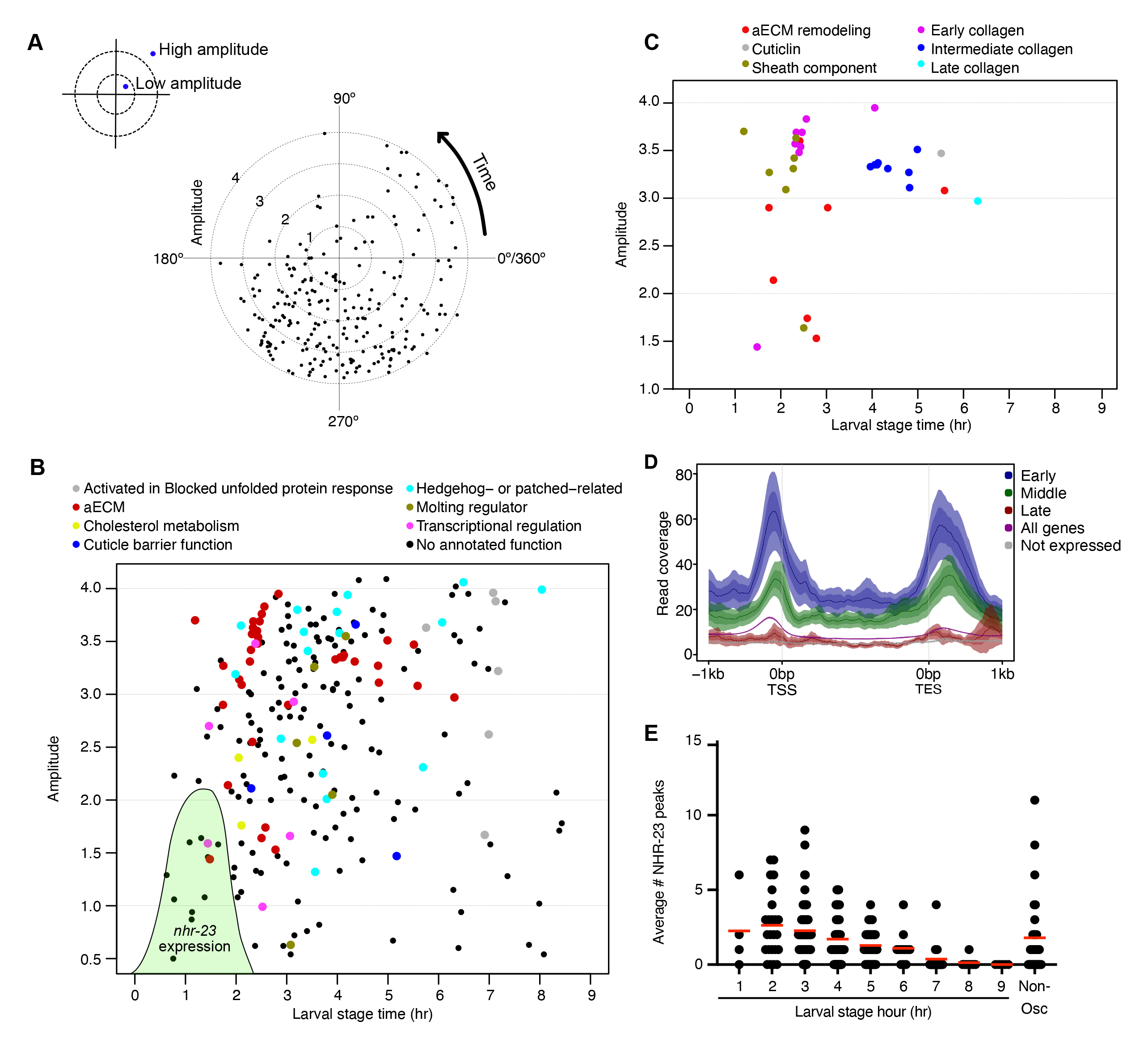
NHR-23 regulates oscillating genes involved in aECM biogenesis. (A) Radar chart plotting amplitude over the phase of peak expression of *nhr-23*-regulated genes from Kouns *et al*. (2011). The dotted circles indicated amplitude, with the innermost circle representing an amplitude of 1 and the outermost circle representing an amplitude of 4. (B) The data from (A) was plotted as a scatter plot and functionally annotated (Table S2). We converted the 360° oscillation to developmental timing over a 9 hour larval stage so that 1 hour=40°. The green shaded area represents NHR-23 expression based on RNA-seq (peak phase=178.11°, amplitude=2.11; Meeuse et al., 2020), imaging (Figure 1A,D,E), and western blotting data (Figure 1B,C). (C) The genes in the aECM component class were subdivided into more specific classes and plotted as in B. (D) Average signal from NHR-23 ChIP-seq data (Gerstein et al., 2010) for *nhr-23*- regulated oscillating genes with peak phases between 135°-254.9° (Early), 255-14.9° (Middle), and 15°-134.9 (Late). *nhr-23* mRNA peaks in expression at 178.11°. Signal is plotted relative to transcription start site (TSS) and transcription end site (TES). The average NHR-23 signal for all 21,600 *C. elegans* genes (All genes) and the 20% of genes with the lowest expression (No expression) are shown for reference. The mean signal is plotted with a line and the mean’s 95% confidence interval is indicated by the shaded area. (E) Number of NHR-23 peaks flanking and within *nhr-23-*regulated genes. The genes are binned by their peak phase relative to adjusted larval stage time. Non-oscillating (Non-Osc) *nhr-23* genes are also depicted.

To explore NHR-23 direct targets, we analyzed an NHR-23 L3 ChIP-seq dataset (Gerstein et al., 2010). As expected, NHR-23 was enriched in the promoter region of genes (Fig. 3D, S5, Table S5). There was also notable enrichment downstream of the transcriptional end site (Fig. 3D, S5, Table S5). We then examined whether NHR-23 was enriched near *nhr-23-*regulated oscillating genes (Table S2). Genes that peaked in expression in the hour following the *nhr-23* mRNA peak in expression (Fig. 3E; hour 2) had the highest average number of NHR-23 peaks flanking and within their gene body. The average number of NHR-23 peaks/promoter declined and few peaks were detected flanking genes between larval stage hours 7-9 (Fig. 3E). Genes which peak in expression close to when *nhr-23* mRNA levels peak tend to have higher NHR-23 levels upstream, downstream, and within their gene bodies (Fig. 3D early genes).

As NHR-23-regulated oscillating genes were enriched in proteases and protease inhibitors (Table S2), we tested how NHR-23 depletion affected expression of a *nas-37* promoter reporter; *nas-37* is a protease implicated in *C. elegans* ecdysis (Davis et al., 2004). We released synchronized *nhr-23::AID, TIR1* L1 larvae carrying a *nas-37p::GFP::PEST* reporter onto control and auxin plates and monitored GFP expression over development. Animals growing on control media displayed oscillating reporter activity with peaks at 18 and 30 hours (Fig. 4A, B). NHR-23 depleted animals exhibited a delayed onset of reporter expression and only a subset of animals expressed the reporter (Fig. 4A, B). There was only one pulse of *nas-37p::GFP::PEST* expression after which we detected no reporter activity (Fig. 4A, B). Together, these data confirm that NHR-23 regulates *nas-37* expression and that following NHR-23 depletion, there is a weaker pulse of target gene expression followed by a failure to express again.

**Fig. 4.**
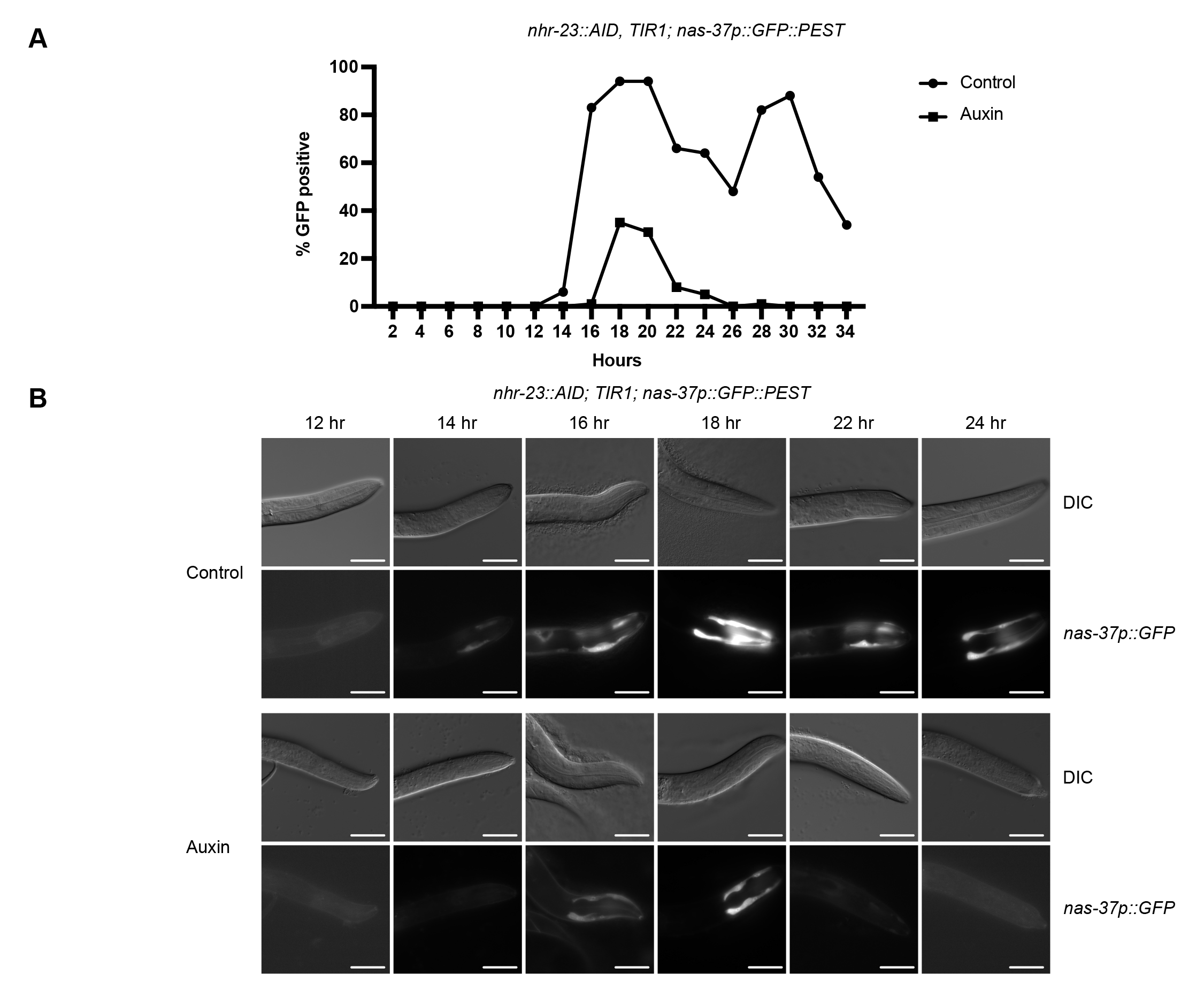
NHR-23-depletion causes reduced expression of a *nas-37p::GFP::PEST* promoter reporter. (A) *nas-37p::GFP::PEST* expression timecourse. Synchronized *nhr-23::AID, TIR1; nas-37p::GFP::PEST* L1 larvae were released on control or auxin plates and scored for head or hypodermal GFP expression every two hours. Fifty animals per time point were scored in two independent experiments and the percentage of animals expressing GFP are presented. (B) Representative images of *nhr-23::AID, TIR1; nas-37p::GFP::PEST* animals grown on control or auxin plates at the indicated time points. Scale bars=20 µm.

### NHR-23 is necessary for NOAH-1 localization to the aECM

Inactivation of the *nhr-23*-regulated predicted protease inhibitor gene *mlt-11* produces embryonic disorganization with accompanying lethality reminiscent of *noah-1* mutants (Ragle et al., 2022; Vuong-Brender et al., 2017). NOAH-1 is a zona pellucida domain protein that is part of the pre-cuticular aECM in embryos and larvae, a transient structure thought to play a role in patterning the cuticle and in embryonic elongation (Cohen and Sundaram, 2020). We introduced an mNeonGreen::3xFLAG (mNG::3xFLAG) tag after the ZP domain which is similar to the insertion site for a previously generated mCherry knock-in (Fig. 5A; Vuong-Brender et al., 2017). *noah-1::mNG::3xFLAG(int)* animals had a wild-type brood size (Fig S6A) and in *noah-1::mNG::3xFLAG, nhr-23::AID; TIR1* lysates there was a band of the expected full length protein size (∼150 kDa) as well as a lower ∼50 kDa band (Fig. 5B). *noah-1* is only predicted to have a single isoform so this might reflect a cleavage or degradation product. ZP proteins are frequently cleaved after their ZP domain (Gupta, 2021; Kiefer and Saling, 2002). While there is a predicted furin RXXR cleavage site at the start of the CFCS domain, the smaller isoform is consistent with cleavage immediately after the ZP domain (Fig. 5A,B). It will be valuable in the future to create isoform-specific mNeonGreen knock-ins to determine where each localizes. NOAH-1::mNG was observed in the cuticle, and in punctate and tubular structures in the hypodermis (Fig 5C). Hypodermal NOAH-1::mNG significantly overlapped with the lysosomal marker NUC-1::mCherry (Fig 5C,D) (Guo et al., 2010; Clancy, Vo et al., 2023). NOAH-1::mNG was detected in lysosomes and weakly in the aECM from L4.1-L4.4 and then reached its maximum aECM intensity between L4.5-L4.7, localizing to thick bands reminiscent of annuli (Fig. 5E). aECM expression decreased from L4.8 to young adulthood (Fig. 5E). NOAH-1::mNG also localized to alae from L4.6 onwards, similar to reports for NOAH-1::sfGFP (Katz et al., 2022).

**Fig. 5.**
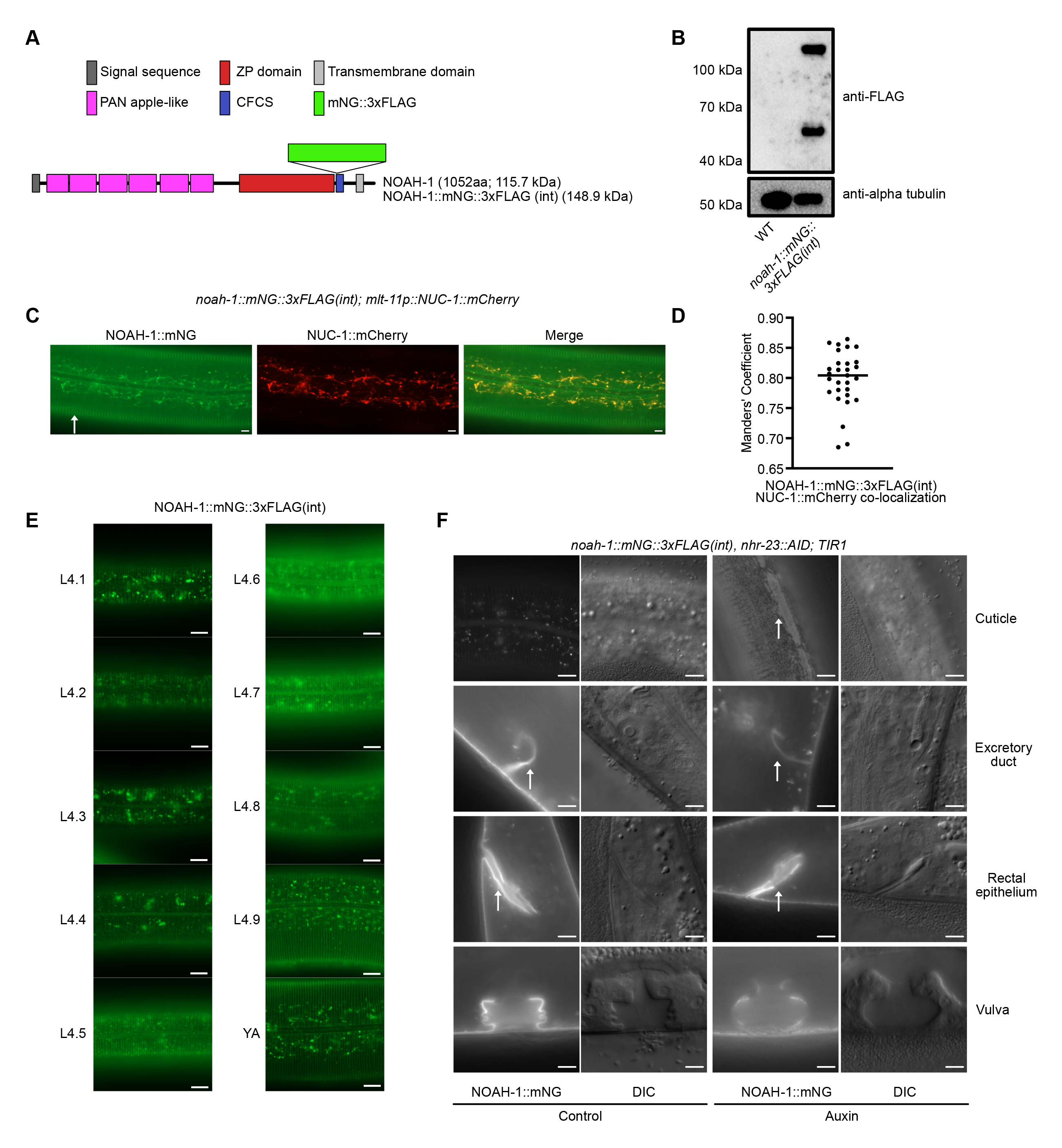
NHR-23-depletion causes defective NOAH-1 localization. (A) Domain prediction for NOAH-1. ZP=Zona Pellucida; CFCS=Consensus Furin Cleavage Site. Predicted size of NOAH-1 and NOAH-1 with an internal mNeonGreen::3xFLAG is provided in kiloDaltons (kDa). (B) Anti-FLAG immunoblots on lysates from *noah-1::mNG::3xFLAG(int)* mid-L4 animals An age-matched WT control is included in each experiment. Marker size (in kDa) is provided and an anti-alpha-tubulin immunoblot was used as a loading control. (C) Representative images from a *noah-1::mNG::3xFLAG(int); mlt-11p::NUC-1::mCherry* mid-L4 animal. Arrow indicates cuticle localization. Scale bars=5 µm. (D) Manders’ co-localization analysis of NOAH-1::mNG::3xFLAG(int) and the lysosome marker NUC-1::mCherry. Three biological replicates were performed analyzing a total of 29 animals. The horizontal line indicates median. (E) NOAH-1::mNG expression timecourse through L4; animals were staged by vulva morphology (Mok et al., 2015). 100 animals were examined over 3 independent replicates. Scale bars=10 µm (F) Representative GFP and DIC images of the indicated tissues of *noah-1::mNG::3xFLAG(int)*, *nhr-23::AID; TIR1,* animals grown on control or auxin plates. NOAH-1::mNG gap over seam cells, and localization to excretory duct and rectum are indicated by arrows. Two biological replicates were performed; the images represent 100% of animals scored (n=51 for control and n=44 for auxin). Scale bars=20 µm for all images except the *noah-1::mNG* excretory duct, rectal epithelium, and vulva (scale bar=5 µm for these images).

We next examined the effect of NHR-23-depletion on NOAH-1 expression and localization. In L4 larvae, we observed the expected vulval lumen localization and aECM expression (Cohen et al., 2020; Katz et al., 2022), as well as localization to the rectum and excretory duct (Fig. 5F). NHR-23 depletion did not affect NOAH-1 localization in the excretory duct or rectum (Fig. 5F). Although vulval morphology is aberrant in NHR-23-depleted animals, NOAH-1 still localizes to cell surfaces and to lysosomes (Fig. 5F). In mid-L4, NOAH-1 is normally localized to the cuticle and lysosomes and is absent over the seam cells (Fig. 5F). NHR-23 depletion caused a loss of NOAH-1::mNG localization to annuli, the formation of small irregular punctae, and accumulation of NOAH-1::mNG over seam cells (Fig. 5F). Together, these data indicate that NHR-23 is dispensable for NOAH-1 expression, but necessary for correct localization in the cuticle.

### NHR-23-depletion causes defects in aECM structure and function

Given the aberrant NOAH-1 localization (Fig. 5) and *nhr-23*-regulation of early and intermediate collagens (Table S2), we next examined whether NHR-23 depletion affected aECM structure. We used CRISPR/Cas9-mediated genome editing in *nhr-23::AID; TIR1* animals to introduce mNG::3xFLAG cassettes into the intermediate collagen *rol-6* (C-terminal translational fusion) and *bli-1,* a collagen that forms struts in the adult cuticle (internal translational fusion)(Fig. 6A). We also introduced a 3xFLAG::mNG into *bli-1* at the same location as the mNG::3xFLAG knock-in. *rol-6* is regulated by *nhr-23* while *bli-1* is an adult-specific collagen expressed in L4 larvae, so its expression would not be interrogated in the microarray experiment (Table S2) (Kouns et al., 2011). We first confirmed that each knock-in had a wild-type brood size (Fig S6A) and then tested fusion protein size by western blotting. Anti-FLAG immunoblotting on *nhr-23::AID; TIR1, rol-6::mNG::3xFLAG* lysates detected a single band of the expected size (∼70 kDA; Fig. 6B). In *nhr-23::AID; TIR1, bli-1::3xFLAG::mNG* there was a single band consistent with N-terminal processing of a monomer at an RXXR furin cleavage site which would remove 10 kDa (125 kDa predicted size). We could only detect protein by extracting soluble cuticle components. BLI-1::3xFLAG::mNG and BLI-1::mNG::3xFLAG had identical localization to one another (Fig. S6B), and were similar to an extrachromosomal transgenic BLI-1::GFP reporter (Tong et al., 2009); all subsequent experiments were performed using *bli-1::mNG::3xFLAG(int)*. Further details of BLI-1::mNG localization will be described elsewhere (Adams et al., submitted).

**Fig. 6.**
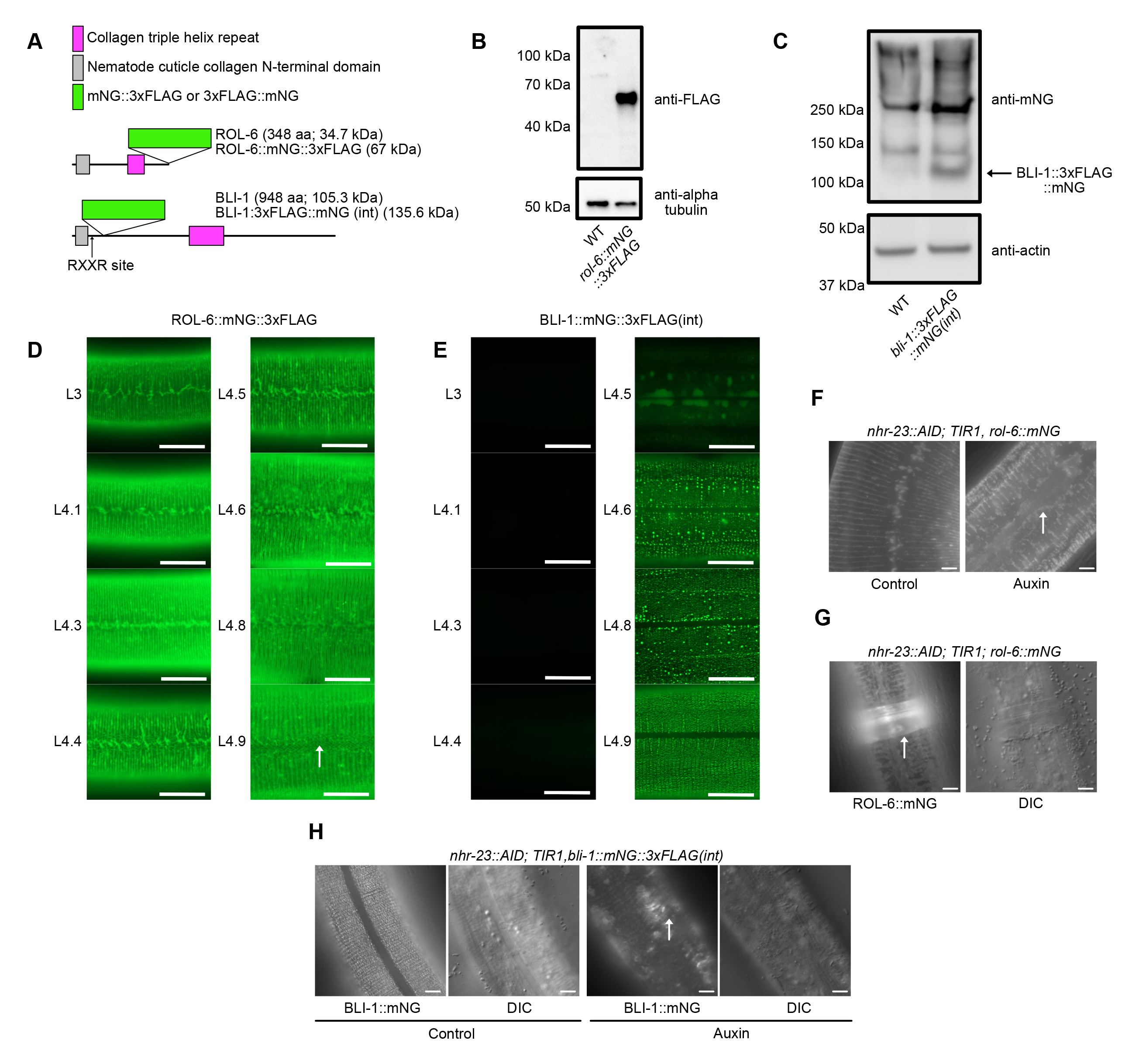
NHR-23-depletion causes ROL-6 and NOAH-1 localization defect and reduced expression and mislocalization of BLI-1. (A) Cartoon of ROL-6 and BLI-1 domains with knock-in position of mNG::3xFLAG or 3xFLAG::mNG tags. Predicted protein size is provided in kilodaltons (kDa). (B) Immunoblots on lysates from *rol-6::mNG::3xFLAG* mid-L4 animals (C) and *bli-1::3xFLAG::mNG(int)* mid-L4 animals (C). An age-matched WT control is included in each experiment. Marker size (in kDa) is provided and an anti-alpha-tubulin (B) or anti-actin (C) immunoblot was used as a loading control. For the blots in (C), the anti-mNeonGreen blot was performed on a soluble cuticle fraction and the anti-actin control was performed on a soluble intracellular fraction (see Methods). An arrow indicates the position of the BLI-1::3xFLAG::mNG band. (D) ROL-6::mNG and (E) BLI-1::mNG expression timecourse through L4; animals were staged by vulva morphology (Mok et al., 2015). 100 animals were examined over 3 independent replicates. Scale bars=10 µm. Arrow indicates fibrous localization pattern in D. (F) Representative images of *nhr-23::AID; TIR1, rol-6::mNeonGreen::3xFLAG* (*rol-6::mNG*) animals grown on control or auxin plates. Arrow indicates gap over seam cells. Images from control and auxin-treated animals are representative of 40/40 animals observed in three independent experiments. (G) Representative images of *nhr-23::AID; TIR1, rol-6::mNG* animals grown on auxin with a corset phenotype. Corset is indicated by arrow. 24/40 animals displayed this phenotype. (H) Representative images of *nhr-23::AID; TIR1, bli-1::mNeonGreen::3xFLAG* (*bli-1::mNG*) animals grown on control or auxin plates. Three experimental replicates were performed. Control images are representative of 42/42 animals scored. For auxin-treated animals, 38/38 exhibited dim expression and 30/38 had patchy BLI-1::mNG localization (indicated by arrow). Scale bars=20 µm in F-H.

We first characterized expression of each translational fusion during the 4^th^ larval stage, using vulval morphology to stage animals (Mok et al., 2015). ROL-6::mNG was first detected in thin bands reminiscent of furrows with a jagged pattern over seam cells (L4.1-L4.3; Fig. 6D). In L4.4-L4.5 some thicker aggregations began appearing and by L4.6-L4.9 ROL-6::mNG re-localized to thicker bands reminiscent of annuli (Fig. 6D). In L4.9, a fibrous pattern could also be observed (Fig. 6D). BLI-1::mNG expression was not detected until L4.5 when it was detected in hypodermal cells (Fig. 6E). In L4.4-L4.8, BLI-1::mNG was observed in a punctate localization in rows with some irregular brighter punctae (Fig. 6E). By L4.9, BLI-1::mNG was found in rows of regularly spaced punctae and was consistently excluded from the cuticle over seam cells throughout L4 and adulthood (Fig. 6E).

To examine the impact of NHR-23 depletion on BLI-1 and ROL-6 localization, we shifted *nhr-23::AID; TIR1; rol-6::mNG* or *nhr-23::AID; TIR1, bli-1::mNG* animals onto control or auxin plates in early L3. We examined animals after 23 hours on control plates when animals were mid-late L4s. We scored NHR-23-depleted animals after 47 hours on auxin plates due to the developmental delay; this approach ensured that control- and auxin-treated animals were stage-matched. ROL-6::mNG expression appeared unaffected by NHR-23-depletion, but the furrow localization was irregular and thicker and we observed gaps in the annular ROL-6::mNG over the seam cells (Fig. 6F). Many animals had a corset of unshed cuticle to which ROL-6::mNG localized (Fig. 6G). We next addressed whether NHR-23 depletion affected the formation of struts in the cuticle medial layer. NHR-23 depletion caused reduced expression of BLI-1::mNG and a loss of the punctate pattern (Fig. 6H). BLI-1::mNG weakly localized to annuli and animals displayed bright, disorganized patches of BLI-1::mNG in the cuticle (Fig. 6H). BLI-1::mNG expression appeared dimmed in the *nhr-23::AID; TIR1* background compared to a wildtype background which would be consistent with low level auxin-independent NHR-23-depletion (Fig S6B).

Given the defects in aECM structure following NHR-23-depletion, we tested whether the epidermal barrier was intact. First, we incubated control and auxin-treated *nhr-23::AID, TIR1* animals with the cuticle impermeable, cell membrane permeable Hoechst 33258 dye which stains nuclei. N2 and *TIR1* animals grown on control or auxin plates and *nhr-23::AID, TIR1* animals grown on control plates exhibited no staining, while almost all animals on auxin plates displayed Hoechst staining (Fig. 7A). Animals shifted onto auxin early or late in L1 robustly expressed an *nlp-29p::GFP* reporter (Fig. 7B, Fig. S7), which can be activated by infection, acute stress, and physical damage to the cuticle (Pujol et al., 2008; Zugasti and Ewbank, 2009). We further tested the cuticle integrity in a hypo-osmotic shock assay in which animals with defective cuticle barriers rapidly die following exposure to water (Kage-Nakadai et al., 2010). *TIR1* animals grown on control media or auxin were completely viable in water (Fig. 7C). *nhr-23::AID, TIR1* control animals had a weakly penetrant sensitivity to hypo-osmotic shock, while the same strain grown on auxin rapidly died following hypo-osmotic shock (Fig. 7C). These data suggest that NHR-23 is required for the cuticle barrier establishment or maintenance.

**Fig. 7.**
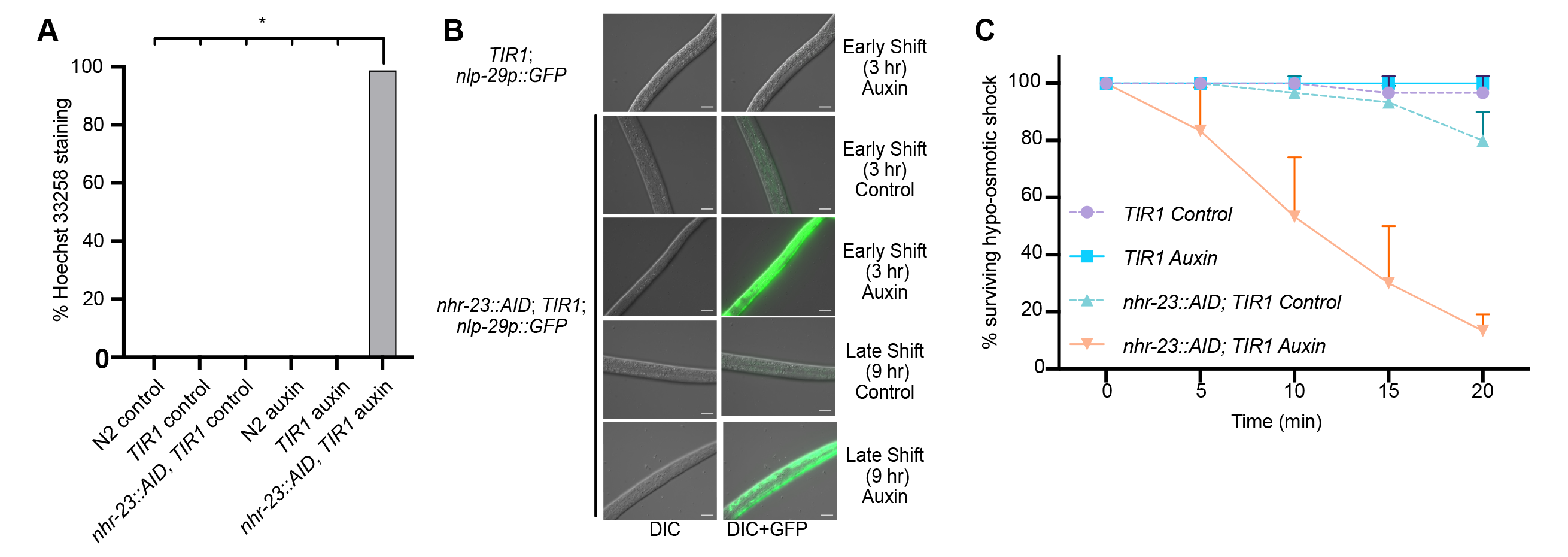
Depletion of NHR-23 leads to a defective cuticle barrier. (A) Permeability to Hoechst 33258 dye. Synchronized animals of the indicated genotype were shifted onto control or auxin plates at 25 hours post-release and grown for 23 hours. Animals were washed off plates and incubated with the cuticle impermeable/membrane permeable Hoechst 33258 dye and scored for nuclear staining in body. The data are from two biological replicates. Asterisk indicates p<0.000001 in two-tailed Student’s t-test. (B) Representative images of GFP expression in animals of the indicated genotype following an early and late shift to control or auxin plates. The images are representative of 50 animals observed during each of two experimental replicates. (C) *nhr-23::AID; TIR1* and *TIR1* animals were subjected to hypo-osmotic shock and scored every five minutes. Ten animals were assayed for each genotype and condition with three biological replicates.

### *nhr-23* is necessary in seam and hypodermal cells for larval development

Finally, we used a tissue-specific RNAi approach to determine in which cells *nhr-23* was necessary for molting. In RNAi-proficient N2 animals, 100% of *nhr-23(RNAi)* animals exhibited developmental delay and molting defects (Fig. 8A). *nhr-23* knockdown in body-wall muscle and intestinal cells produced no molting defects and we observed only mild developmental delay (Fig. 8A). Vulval precursor cell-specific *nhr-23(RNAi)* caused moderate molting defects and developmental delay (Fig. 8A). We observed highly penetrant developmental delay and molting defects when we performed *nhr-23* RNAi in a strain using a *wrt-2* promoter for tissue-specific RNAi (Fig. 8A). *wrt-2* is expressed in seam cells, rectal cells and hypodermal cells (Fig. S8A, B; Aspöck et al., 1999). Given this result and the defective ROL-6::mNG and NOAH-1::mNG (Fig. 5,6) localization over seam cells, we generated more tissue-restricted RNAi strains. We used a minimal seam cell-specific enhancer with a *pes-10* minimal promoter (Ashley et al., 2021), which is robustly expressed in seam cells with some weak hypodermal expression (Fig. S8C, D). To test the specificity of this strain, we introduced a *his-72::mNG* knock-in. In animals treated with control RNAi, we observed nuclear expression in seam, hypodermal, intestinal, vulval and germline cells, as expected for a histone H3 fusion (Fig S9). *mNeonGreen* RNAi caused reduced HIS-72::mNG expression in seam, hypodermal syncytium, and intestinal nuclei. *nhr-23(RNAi)* in this *SCMp* RNAi strain phenocopied *nhr-23(RNAi*) in N2 animals with almost all animals exhibiting developmental delay and molting defects (Fig. 8A). Notably, intestine-specific *nhr-23* RNAi did not cause phenotypes (Fig. 8A). We then constructed a hypodermis-specific RNAi strain using the *semo-1* promoter (Kaletsky et al., 2018; Köhnlein et al., 2020). *nhr-23* RNAi in this strain produced molting defects and developmental delay, though with less penetrance compared to our *SCMp-*specific RNAi strain (Fig 8A). To test the effect of *SCMp*- specific *nhr-23* knockdown on the aECM, we introduced a BLI-1::mNG knock-ins in our seam cell-specific RNAi strain. *nhr-23* depletion in this strain caused reduced BLI-1 levels relative to control RNAi (Fig. 8B). This BLI-1::mNG reduction was comparable to both *nhr-23* RNAi in a wild-type control (Fig. 8B) and to NHR-23 protein depletion in the soma (Fig. 6H). Together these data indicated that *nhr-23* is necessary in the seam and hypodermal cells for timely developmental progression, completion of molting, and aECM formation.

**Fig. 8.**
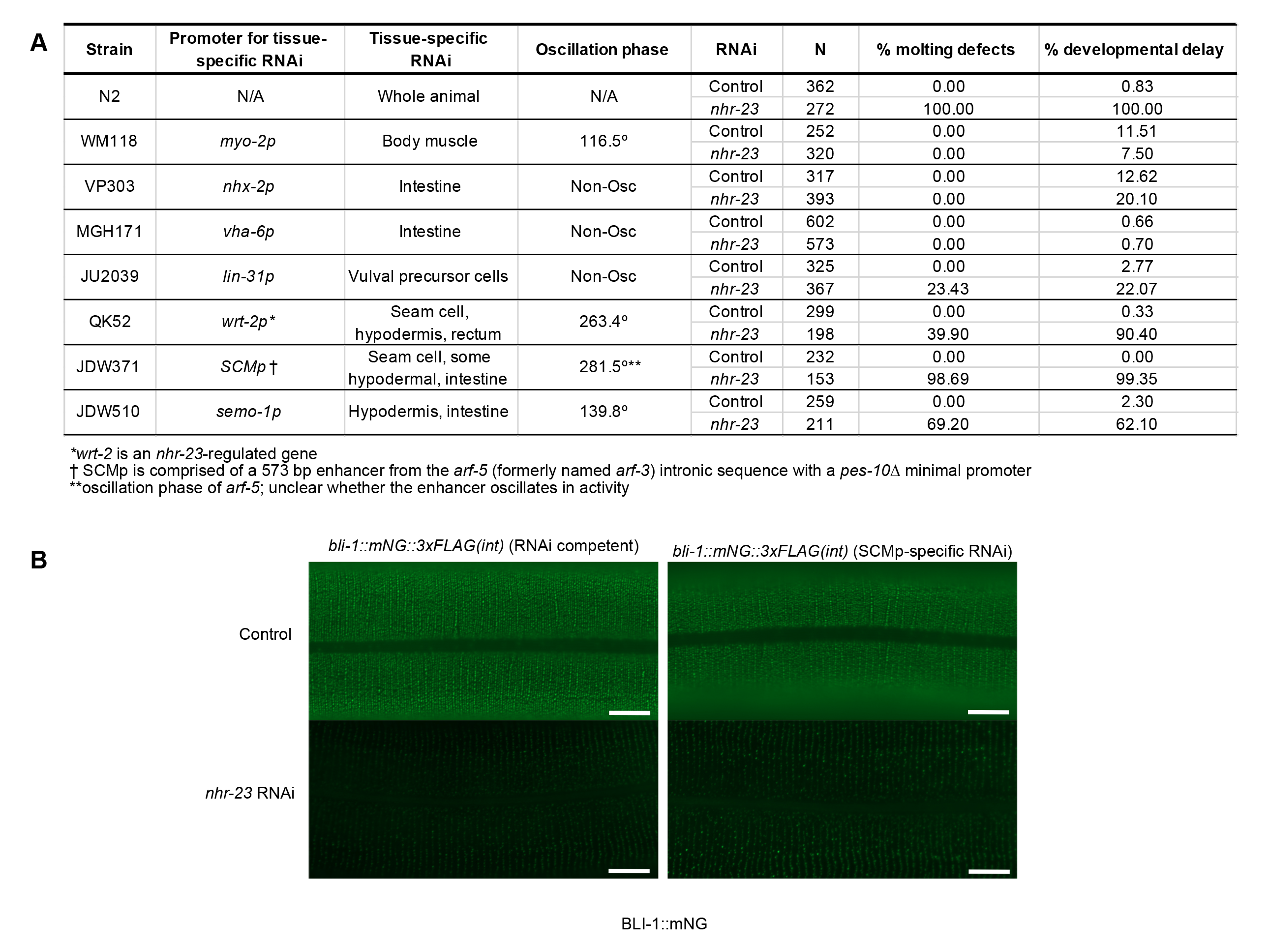
*nhr-23* is necessary in seam cells for developmental progression, molting and BLI-1 localization. (A) Tissue-specific RNAi. A timed egg lay of animals of the indicated strains was performed on control or *nhr-23* RNAi plates and plates were scored three days later. The promoters used to reconstitute RNAi, their tissue-specificity, and oscillation phase (when applicable) is provided. The number of progeny scored from two biological replicates is indicated. Developmental delay was scored as a failure to reach adulthood after 72 hours of growth. Molting defects included animals trapped in cuticles, animals dragging cuticles, and animals with cuticle corsets. (B) L2 *bli-1::mNG* (RNAi competent) and *bli-1::mNG; SCMp::rde-1; rde-1(ne300)* (*SCMp*- specific RNAi) animals were and shifted onto control or *nhr-23* RNAi plates at L2 and grown until stage L4.8. Equal exposure times were used for all images; scale bars=10 µm. Images are representative of 20 animals over two independent experimental replicates.

## DISCUSSION

*C. elegans* molting is a powerful system to understand developmentally programmed aECM regeneration. In this study, we determine when and where NHR-23 acts to promote molting. NHR-23 oscillates and its depletion causes severe developmental delay. *nhr-23*-regulated genes are enriched for cuticle components and protease inhibitors, and NHR-23 is enriched at the transcription start and end sites of target genes. NHR-23 depletion causes aberrant localization of the early collagen ROL-6 and the pre-cuticle component, NOAH-1. These cuticle defects are correlated with a loss of the cuticle permeability barrier function. Loss of NHR-23 function in L4 stages also causes severely reduced levels of the adult collagen BLI-1, which is also mislocalized. These cuticle defects are correlated with a loss of the cuticle barrier. Tissue-specific RNAi suggests that *nhr-23* is necessary in seam and hypodermal cells.

### NHR-23 is necessary for cuticle structure and function

NHR-23 binds more robustly in the promoter and transcription end site of genes which peak in expression closer to the *nhr-23* mRNA peak in expression (Fig. 3). One intriguing possibility is that the earlier NHR-23-regulated genes may be more sensitive to NHR-23 levels. Consistent with this model, there appear to be more NHR-23 peaks flanking and within early *nhr-23-*regulated genes (Fig. 3, Tables S2, S5). This model would align with *E. coli* amino acid biosynthesis gene regulation where enzymes earlier in the pathway have more responsive promoters with higher activity (Zaslaver et al., 2004). Interestingly, the non-oscillating *nhr-23-*regulated genes had a range in number of NHR-23 peaks. It’s unclear why NHR-23 oscillation is failing to drive oscillation of these target genes. Some possible explanations include that these genes could have stable mRNA transcripts in comparison to the oscillating genes or there is combinatorial gene regulation involving a non-oscillating transcription factor.

NHR-23 regulates the expression of *nas-37*, a protease implicated in molting (Fig. 4; Frand et al., 2005). Curiously, two other targets (*noah-1, rol-6*) seemed to be expressed at near wild-type levels but displayed aberrant localization (Fig. 5,6). These data highlight a potential limitation of a single timepoint gene expression study. A shift in phase of an oscillating gene without change in amplitude could create the appearance of up- or down-regulation depending on the timepoint sampled (Tsiairis and Großhans, 2021). The *nhr-23(RNAi)* microarray is a useful starting point to understand how NHR-23 promotes molting, but an RNA-seq timecourse on NHR-23 depleted animals may be necessary to identify regulated genes. Such an approach was necessary to determine how the pioneer factor, BLMP-1, promotes developmental timing (Hauser et al., 2021).

NHR-23-depletion caused aberrant localization of NOAH-1 and ROL-6, lower levels of BLI-1, and a severe barrier defect (Fig. 5,6,7). There are numerous *nhr-23*-regulated genes which are implicated in the epithelial barrier. The cuticle furrow formed by six *nhr-23*-regulated collagens (*dpy-2, dpy-3, dpy-7, dpy-8, dpy-9, dpy-10*) is thought to be monitored by a sensor which coordinates several stress responses (Dodd et al., 2018). An RNAi screen of 91 collagens found that inactivation of only these six collagens caused a barrier defect (Sandhu et al., 2021). Inactivation of a subset of the furrow collagens causes permeability to Hoechst 33458 dye and elevated *nlp-29p::GFP* reporter activity (Dodd et al., 2018), consistent with our NHR-23-depletion data (Fig. 7). *bus-8* is a predicted glycosyltransferase that plays a role in the epithelial barrier and is also *nhr-23*-regulated and its peak expression follows that of the furrow collagens (Table S2). Screening *nhr-23* regulated genes may reveal other genes implicated in the epithelial barrier and the peak phase could provide insight into how this barrier is constructed.

### NHR-23-depletion causes developmental delay and failed molting

What is driving NHR-23-depleted animals to eventually attempt to molt? An NHR-23- regulated promoter reporter (*nas-37p::GFP::PEST*) peaked at a similar time in control and NHR-23-depleted animals (Fig. 4), but was expressed at lower levels. This data might suggest that the low levels of NHR-23 present are sufficient to drive low levels of target gene expression. Consistent with this idea, NHR-23 depletion or knockdown results in developmental delay but not an arrest (Fig. 2; MacNeil et al., 2013; Patel et al., 2022). The pre-cuticle component, NOAH-1, and early collagen, ROL-6 are both expressed and secreted but display localization defects. One possibility is that low sustained NHR-23-levels eventually allow the accumulation of factors that initiate apolysis but there is insufficient expression for execution. *nas-37* is necessary for *C. elegans* ecdysis and recombinant NAS-37 promotes the formation of retractile rings, a structure associated with molting, in the parasitic nematode *H. contortus* (Davis et al., 2004). Another non-exclusionary possibility is that NHR-23 controls the expression of transcription factors in a regulatory cascade, like the gene regulatory network controlling insect molting (King-Jones and Thummel, 2005; Thummel, 1990). *nhr-23* regulates *peb-1* and *dpy-20*, transcription factors with known roles in molting (Clark et al., 1995, 20; Fernandez et al., 2004, 1). *nhr-23* also regulates *nhr-91* (Kouns et al., 2011)(Table S2), a nuclear hormone receptor involved in blast cell progression in response to nutrients (Kasuga et al., 2013). A third possibility is that another timer eventually promotes molting. The molting timer and heterochronic cycles can be uncoupled, causing animals to attempt molting before completion of cell division and differentiation, leading to death (Ruaud and Bessereau, 2006).

### *nhr-23* is necessary in seam and hypodermal cells to promote molting

Seam cell-enriched *nhr-23* RNAi caused severe developmental delay and molting defects (Fig 8). Hypodermal-enriched *nhr-23* knock-down produced less penetrant phenotypes (Fig. 8). The most severe defects in ROL-6 and NOAH-1 localization occur above the seam cells (Figure 6). Determining how NHR-23 regulates BLI-1 expression and formation of struts will provide insight into aECM assembly. *nhr-23* is expressed in other epidermal cells that produce the pre-cuticle and cuticle, such as vulval precursor cells, rectal cells, and excretory duct cells. NHR-23 depletion did not produce obvious defects in localization of the pre-cuticle component NOAH-1 in these cells (Fig. 5). We did not see phenotypes such as excretory duct or pore lumen dilation, which would be indicative of excretory system defects (Gill et al., 2016). We observed vulval morphology defects following NHR-23 depletion, but it is not clear whether that is due to NHR-23 regulation of this specialized aECM or if it is the consequence of developmental delay and failure to molt. Tissue-specific RNAi or protein depletion will be required to determine whether NHR-23 activity is necessary in these epithelial cells, or if it predominantly functions in seam and hypodermal cells. Our data indicate the importance of tissue-specific RNAi strain validation. *SCMp* drove reporter expression robustly in seam cells (Fig. S8). However, when we made a tissue-specific RNAi strain using *SCMp* we observed depletion in seam, hypodermal, and intestinal cells (Fig. S9). RNAi has an inherent amplification of dsRNA triggers by RNA-dependent RNA polymerases (Zhang and Ruvkun, 2012). Weak promoter activity in undesired tissues could reconstitute RNAi in these tissues. A systematic analysis of tissue-specific RNAi strains is an important future direction for interpreting studies using these strains.

### Future perspectives

This work highlights the power of timed protein depletion for dissecting the role of oscillating developmental regulators in development. As transcription factors coordinate the expression of batteries of genes in a given biological process, future work will reveal how NHR-23 coordinates apical ECM remodeling, apolysis, and ecdysis.

## MATERIALS AND METHODS

### *C. elegans* Strains and Culture

*C. elegans* strains were cultured as originally described (Brenner, 1974), except worms were grown on MYOB instead of NGM. MYOB was made as previously described (Church et al., 1995). Animals were cultured at 20°C for all assays, unless otherwise indicated. For general strain propagation, animals were grown at 15°C according to standard protocols. Brood sizes were performed picking L4 larvae to individual wells of a 6-well plate seeded with OP50 and incubating the plate at 20°C. Animals were transferred to new plates daily over 4 days. Two days post-transfer, the number of hatched progeny and unfertilized eggs were scored.

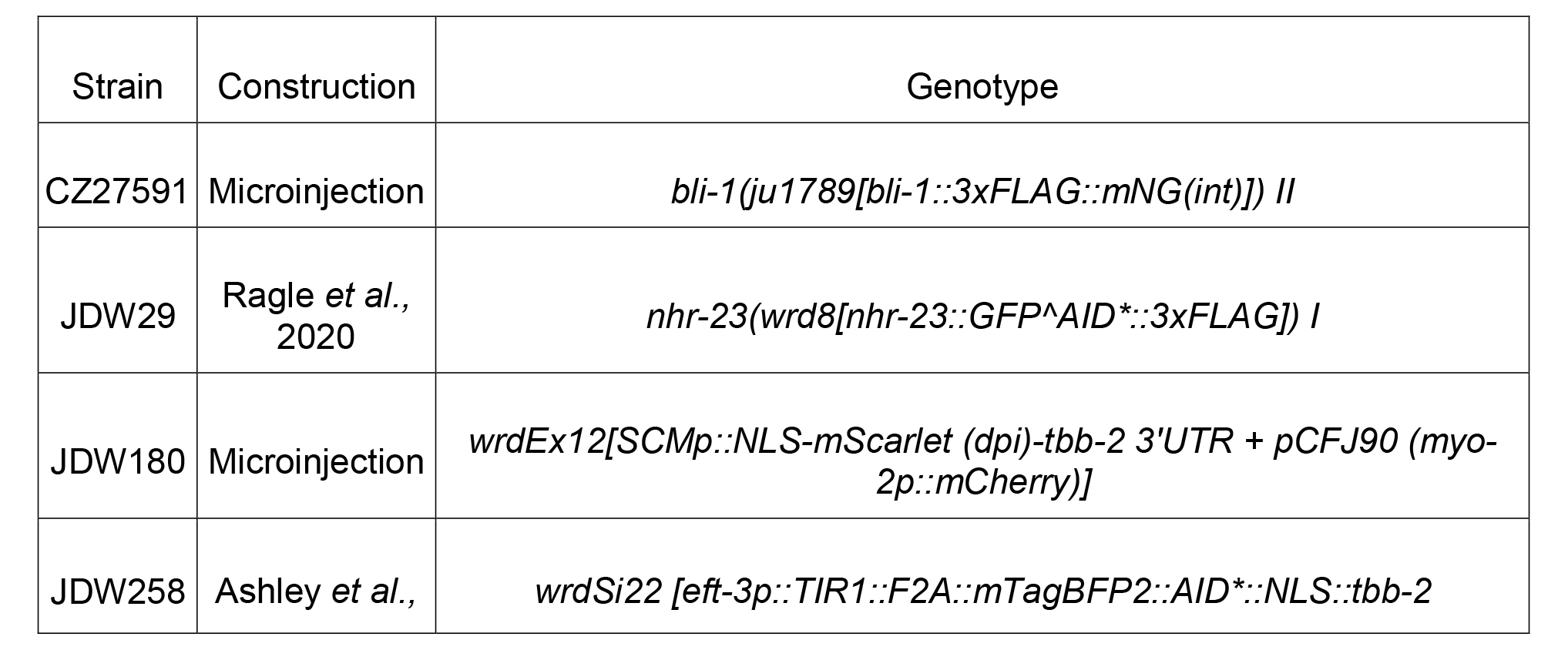

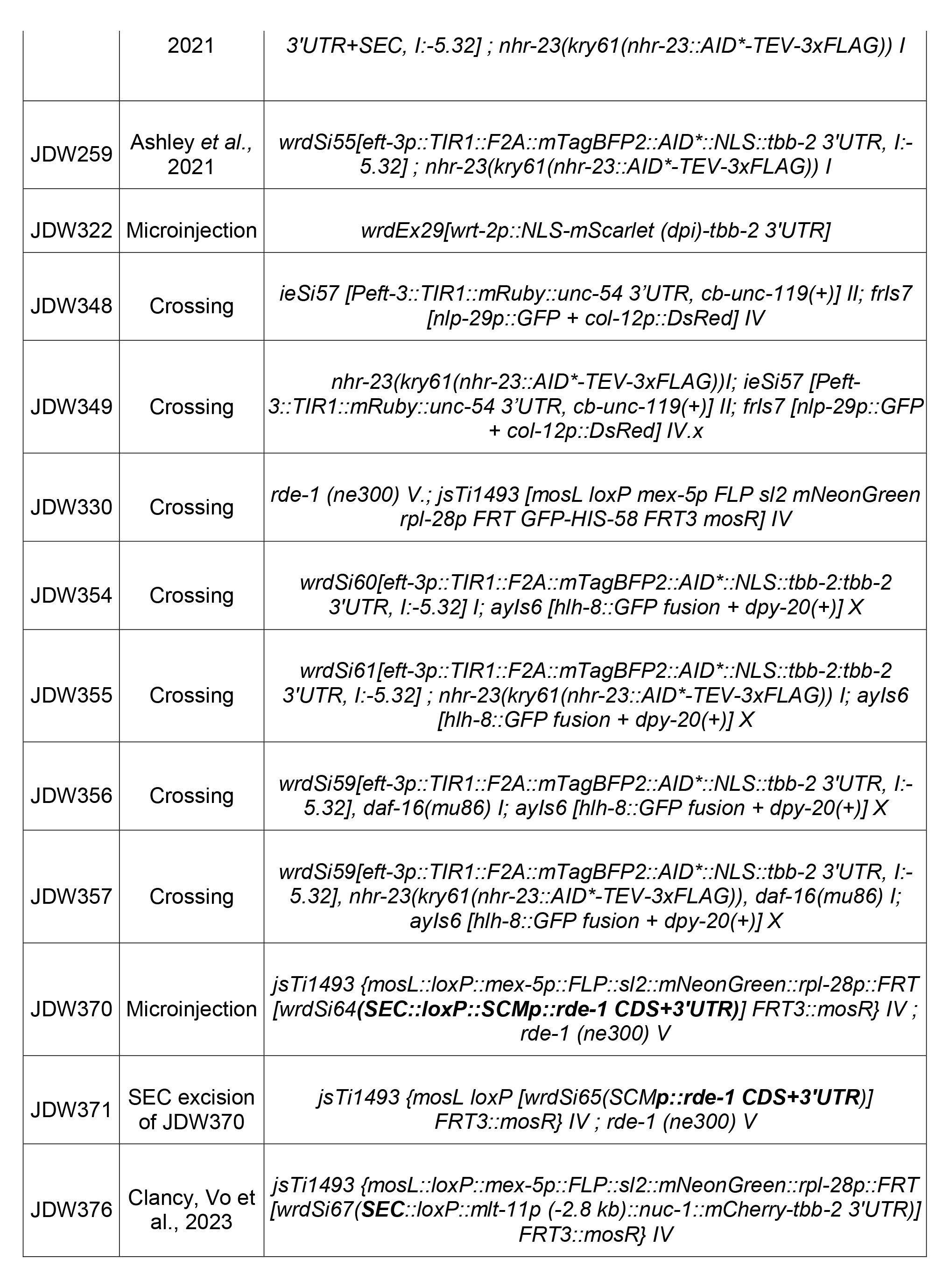

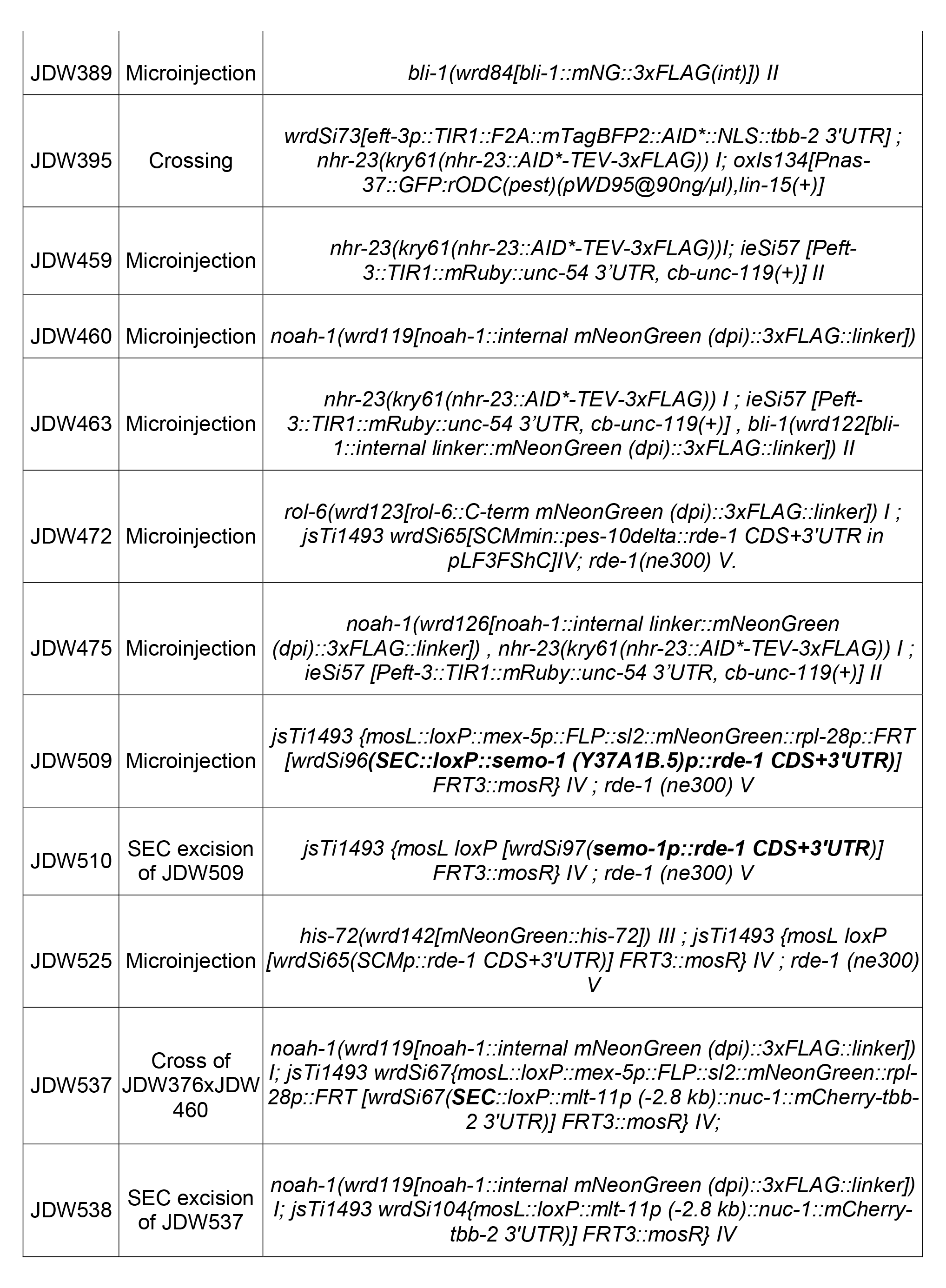

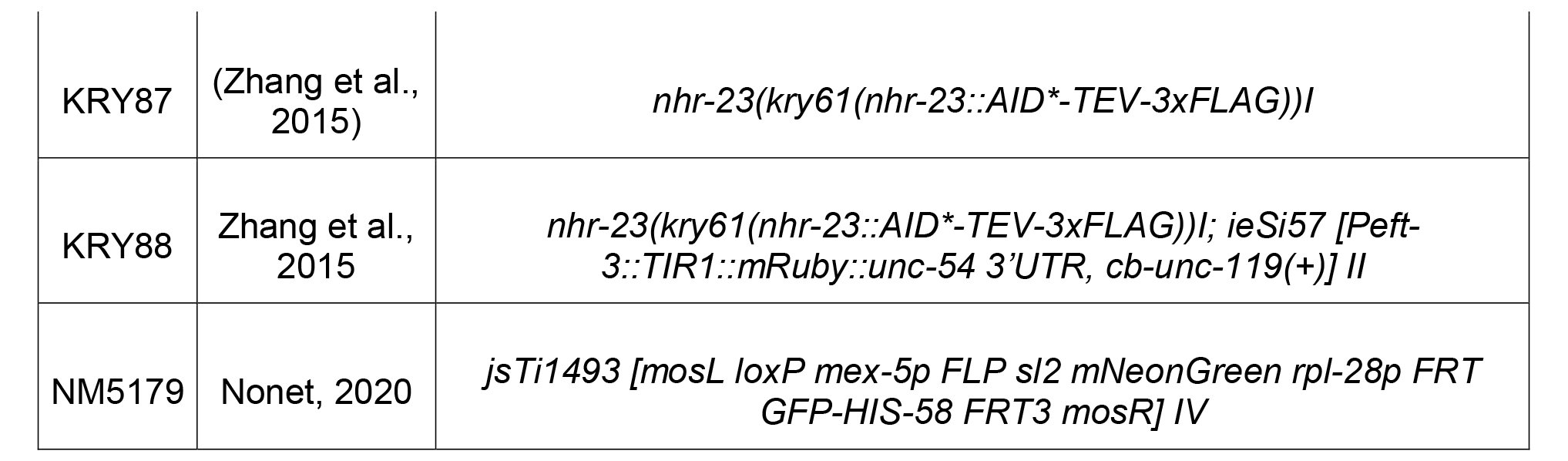
Strains used in this study:

Strains provided by the *Caenorhabditis* Genetics Center:

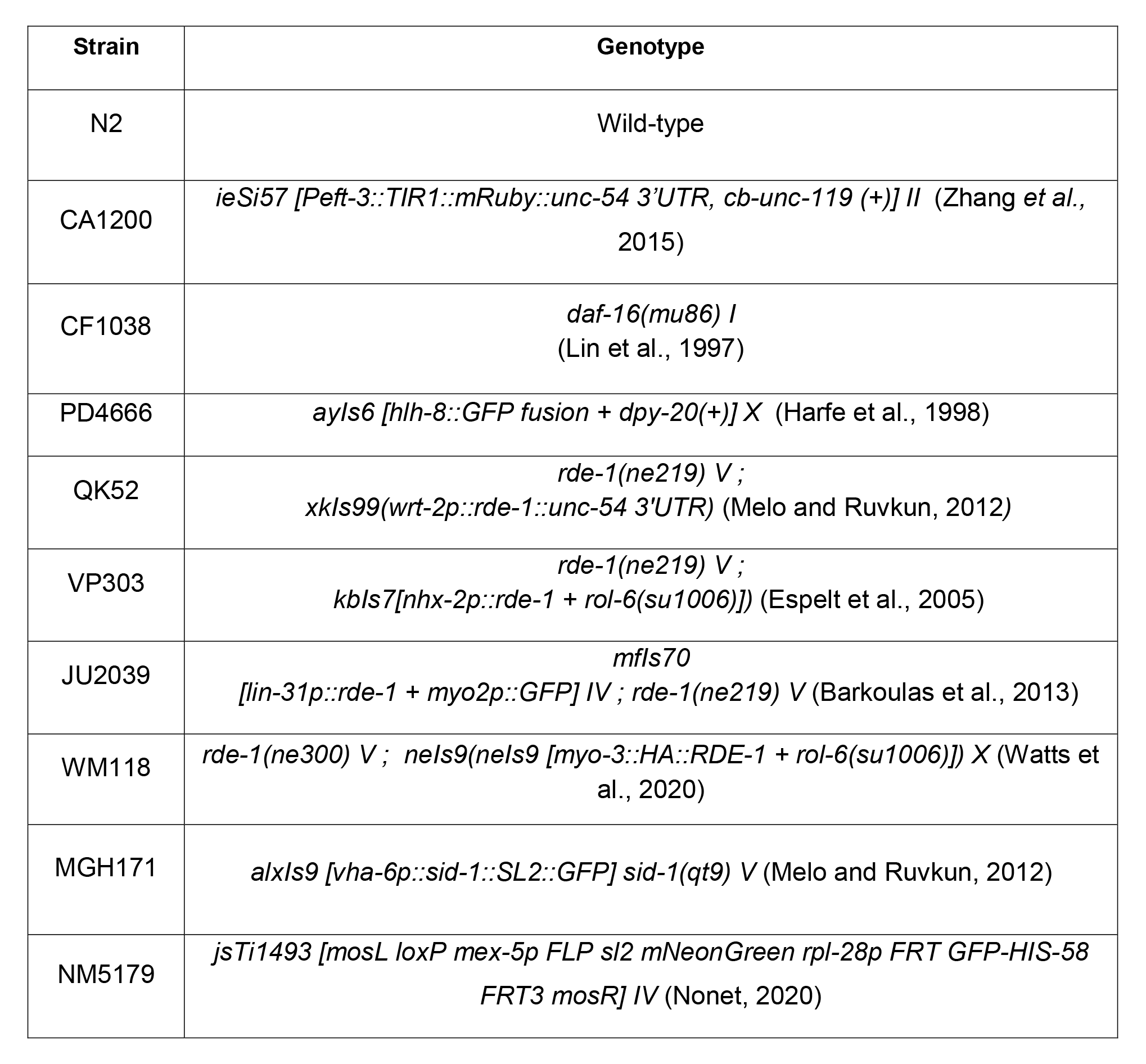

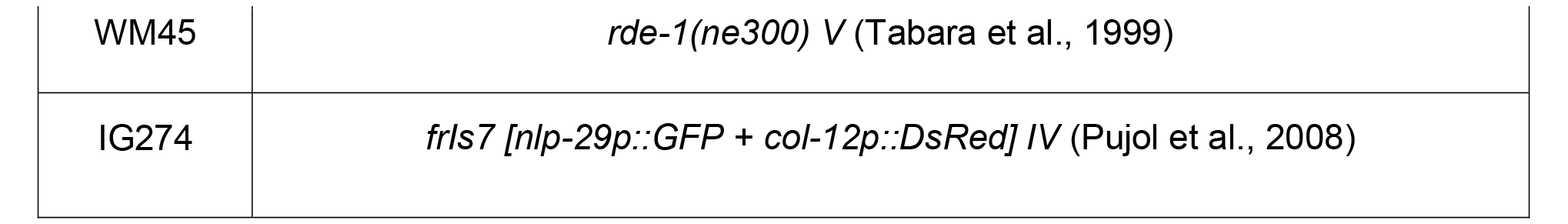

### Other strains

EG3200 oxIs134[Pnas-37::GFP:rODC (PEST) (pWD95@90ng/µl),lin-15(+)]; lin-15(n765ts) X (Davis et al., 2004) was provided by D. Fay (U Wyoming).

### Genome editing and transgenesis

*mNeonGreen::3xFLAG* knock-ins into *bli-1, noah-1,* and *rol-6* and *mNeonGreen* knock-ins into *his-72* were generated by injection of Cas9 ribonucleoprotein complexes [700 ng/µl IDT Cas9, 115 ng/µl crRNA and 250 ng/µl IDT tracrRNA] and a dsDNA repair template (25-50 ng/ul) created by PCR amplification of a plasmid template (Paix et al., 2014; Paix et al., 2015). The *bli-1* and *noah-1* knock-ins were internal and the *mNG::3xFLAG* cassette was flanked by flexible glycine- and serine-rich linker sequences. Generation of the *BLI-1::3xFLAG::mNG(int)* used in Fig 6C will be described elsewhere (Adams et al submitted). The C-terminal *rol-6::mNG::3xFLAG* fusion encoded a flexible linker between the end of *rol-6* and the start of mNG. The *noah-1::mNG::3xFLAG(int)* knock-in used a pJW2332 repair template, which will be described elsewhere; details available upon request. The PCR products were melted to boost editing efficiency, as previously described (Ghanta and Mello, 2020). crRNAs used are provided in Table S6. Oligonucleotides used for repair template generation from template pJW2172 (Ashley et al., 2021) and for genotyping are provided in Table S7. Genomic and knock-in sequences are provided in File S1. Plasmids used are provided in Table S8. To generate JDW371 and JDW510 (seam cell and hypodermal-specific tissue-specific RNAi) we crossed a *jsTi1493* landing pad for recombination-mediated cassette exchange (RMCE)(Nonet, 2020) into an *rde-1(ne300*) null mutant, generating JDW330. We used Gibson cloning to introduce a promoterless *rde-1* genomic coding sequence+3’UTR fragment into the RMCE integration vector pLF3FShC (Nonet, 2020), creating pJW2247. This vector can be linearized with *Avr*II+*Bsi*WI double-digestion and promoters can be introduced by Gibson cloning (Gibson et al., 2009). We generated a *SCMp* (seam cell-specific) and *semo-1p* (hypodermis specific) by Gibson cloning promoters into linearized pJW2247, constructing pJW2236 and pJW2264, respectively. These vectors were integrated into *jsTi1493* landing pad in JDW330 using RMCE, as described (Nonet, 2020). *wrt-2p* and *SCMp* promoter reporters were constructed in pJW1836 (*NLS::mScarlet::tbb-2 3’ UTR*)(Ashley et al., 2021) by Gibson cloning. Plasmids were injected into N2 animals at 50 ng/µl with a pCFJ90 co-injection marker at 10 ng/µl for the *SCMp* promoter reporter and a pRF4 co-injection marker at 50 ng/µl for the *wrt-2* promoter reporter (Frøkjaer-Jensen et al., 2008; Mello et al., 1991). Transgenic lines propagating extrachromosomal arrays were generated as previously described (Mello et al., 1991). Sequence files for plasmids are provided in File S2. Primers and cloning details are available upon request.

### Auxin treatment

Control and auxin media/plates were made as described in Ragle et al. 2020. Control media consisted of MYOB agar+0.25% ethanol. Auxin media was made by dissolving indole 3-acetic acid (IAA; Alfa Aesar, AAA1055622) in 100% ethanol to 1.6 M and then mixing it into melted MYOB agar at 55°C to a final concentration of 4 mM prior to pouring plates. Control plates contained 0.25 % ethanol. Temperature of the media was monitored with a Lasergrip 1080 infrared thermometer gun (Etekcity). Plates were seeded with *E. coli* OP50 and incubated overnight at room temperature. Plates were stored for up to one month at 4°C prior to use. For most auxin treatment experiments, animals were synchronized by alkaline bleaching (dx.doi.org/10.17504/protocols.io.j8nlkkyxdl5r/v1). The collected eggs were incubated in M9 buffer supplemented with 5 mg/mL cholesterol at 20°C for 24 hours and arrested L1 larvae were released onto the indicated type of MYOB plate. For the experiment in Table S1 a timed egg lay was performed. Twenty adult hermaphrodites animals of the indicated genotype were picked onto control (0.25% ethanol) or auxin (4 mM IAA) plates and allowed to lay eggs for 2 hours. The adult animals were removed and plates were incubated for 48 hours at 25°C.

### Microscopy

Synchronized animals were collected from MYOB, control, or auxin plates by either picking or washing off plates. For washing, 1000 µl of M9 + 0.05% gelatin was added to the plate or well, agitated to suspend animals in M9+gelatin, and then transferred to a 1.5 ml tube. Animals were spun at 700xg for 1 min. The media was then aspirated off and animals were resuspended in 500µl M9 + 0.05% gelatin with 5 mM levamisole. 12 µl of animals in M9 + 0.05% with levamisole solution were placed on slides with a 2% agarose pad and secured with a coverslip. For picking, animals were transferred to a 10 µl drop of M9+5 mM levamisole on a 2% agarose pad on a slide and secured with a coverslip. Images were acquired using a Plan-Apochromat 40x/1.3 Oil DIC lens, a Plan-Apochromat 63x/1.4 Oil DIC lens, or a Plan-Apochromat 100x/1.4 OIL DIC lens on an AxioImager M2 microscope (Carl Zeiss Microscopy, LLC) equipped with a Colibri 7 LED light source and an Axiocam 506 mono camera. Acquired images were processed through Fiji software (version: 2.0.0- rc-69/1.52p) (Schindelin et al. 2012). For direct comparisons within a figure, we set the exposure conditions to avoid pixel saturation of the brightest sample and kept equivalent exposure for imaging of the other samples. Co-localization using a Manders’ co-efficient (Manders et al., 1992; Manders et al. 1993) was performed as described in Clancy, Vo et al., 2023.

### Phenotypic analysis

For the phenotypic analysis in Figure 2, synchronized CA1200 or KRY88 larvae were released onto MYOB plates and then shifted onto control or auxin plates every two hours up to 16 hours. Animals were collected as described in the Microscopy section and imaged by DIC microscopy to score for morphology and shedding of the L1 cuticles. For the M cell lineage experiments in Figure S3, synchronized animals of the indicated genotypes were released onto MYOB plates and shifted onto control or auxin plates at 3 hours (early shift) or 9 hours (late shift) post-release and then imaged at 24 hours post-release, as described in the Microscopy section. M cells were counted and recorded. For the analyses in Figure 2E, synchronized *nhr-23::AID, TIR1* larvae were released on auxin plates and scored for viability and molting defects 72 hours later. For the L3 shift experiments (Fig. 2F,G) synchronized animals of the indicated genotype were grown on MYOB. At 25 hours post-release, they were shifted onto 6-well control or auxin MYOB plates seeded with OP50. Animals were collected 23 or 72 hours later as described in the Microscopy section and imaged by DIC microscopy using a 63x lens to stage animals according to vulval morphology (Mok et al., 2015). For the reporter timecourse, synchronized *nhr-23::AID, TIR1, nas-37p::GFP::PEST* animals were released on control or auxin plates. They were then scored for GFP expression using a PlanApo 5.0x/0.5 objective on a M165 FC stereomicroscope (Leica) equipped with an X-cite FIRE LED lightsource (Excelitas) and long pass GFP filter set (Leica #10447407). We scored GFP expression in the head and hypodermis and did not score rectal GFP expression, as expression perdured in this tissue after head and hypodermal GFP expression ceased.

### Barrier assays

Hoechst 33258 staining was performed as described previously (Moribe et al., 2004), except that we used 10 µg/ml of Hoechst 33258 as in Ward et al. (2014). Two biological replicates were performed examining 50 animals per experiment. The number of animals with either head or hypodermal nuclei staining with Hoechst 33258 were scored under a 10x DIC objective. Representative images were then taken with equivalent exposures using a 63x Oil DIC lens, as described in the imaging section. Hypo-osmotic stress sensitivity assays were performed on L4 stage larvae as described (Ward et al., 2014), except we used 20 µl of dH_2_0. Each strain was assayed in triplicate.

### Western Blots

Animals were synchronized by alkaline bleaching, as described in the “Auxin Treatment” section. For the blots in Fig. 5B and 6B, 30 animals of the indicated genotype and stage were picked into 30 µl of M9+0.05% gelatin and 10 µl of 4x Laemmli sample buffer was added and then samples were heated at 95°C for 5 minutes and stored at −80°C until they were resolved by SDS-PAGE. For the blots in Fig. 6C, synchronized animals of the indicated genotype were collected at the L4 stage. Extracts of soluble cuticle proteins were prepared as described (Cox *et al*., 1981) with minor modifications. Briefly, worms were collected using M9J buffer in a 15 ml Falcon tube and washed 3X with M9J buffer.

Worm pellets were resuspended in 5 ml of sonication buffer and incubated on ice for 10 min, sonicated (Misonix XL-2000 Series Ultrasonic Liquid Processor, 10 × 10 s pulse with 10 s break) with 30 μl of 0.1 M PMSF. Following sonication, samples were centrifuged at 8000 rpm for 2 min at 4°C and supernatants stored as Fraction 1 (F1; intracellular proteins) at −20°C for the purpose of using it as loading control. Pellets were washed 3X with sonication buffer and the resultant pellets were resuspended in 100 μl sonication buffer, boiled at 95°C for 2 min with 1 ml of ST buffer and incubated overnight with rotation at RT. Following incubation, samples were centrifuged at 8000 rpm for 3 min and supernatants were stored as F2 at −20°C. Subsequently, cuticle pellets were washed 3X with 0.5 % triton X-100 and boiled at 95°C for 2 min with 300 μl of ST buffer with 5% β-mercaptoethanol and incubated with rotation at room temperature for 16 h. Finally, the samples were centrifuged at 8000 rpm for 3 min and supernatants collected and stored as F3 (soluble cuticle proteins). F3 Fraction was precipitated with methanol/chloroform and resuspended in 100 μl of rehydration buffer.

For the remaining blots, animals of the indicated genotype were washed out of wells on a 6-well plate at the indicated timepoints with M9+0.05% gelatin (VWR, 97062-620), transferred to a 1.5 ml tube and washed twice more with M9+0.05% gelatin, as previously described (Vo et al., 2021). Animals were pelleted, transferred in a 100 µl volume to a new 1.5 ml tube, and flash frozen in liquid nitrogen, then stored at −80°C. Prior to SDS-PAGE, samples were freeze-cracked twice in liquid nitrogen or on dry ice, Laemmli sample buffer was added to 1X and samples were heated to 95°C for 5 minutes. For the western blots in Figure 1C and 1D 6000 KRY88 animals per well were transferred onto a 6-well MYOB plate seeded with OP50. For the western blot in Figure 2B, 6000 synchronized KRY88 animals were plated onto control or auxin plates. Control animals were harvested at peak NHR-23 expression (12 hours of growth on control plates) and auxin-treated animals were shifted onto auxin at this time and samples were taken at the indicated time points, as described above. For the western blots in Figure S7, 500 animals of the indicated genotype were transferred into wells of a 6-well control or auxin plate seeded with OP50. Animals were incubated for 24 hours at 20°C and collected as described above.

For the blots in Figs. 1C,D, 10 µl of lysate and 7.5 µl 1:1 mix of Amersham ECL Rainbow Molecular Weight Markers (95040-114) and Precision Plus Protein Unstained Standards (#1610363) was resolved by SDS-PAGE using precast 4-20% MiniProtean TGX Stain Free Gels (Bio-Rad). The blot in Fig. S7, the lysates had more variable levels of total protein, particularly the NHR-23 depleted samples that produced an early larval arrest. Therefore, we quantified the signal of the most intense band by Stain Free imaging (Posch et al., 2013) using a Bio-Rad ChemiDoc imaging system, and ran a new gel with normalized loading. The lysates in Fig. 2A, 5B, 6B, and S7 were resolved as described above except we used a Spectra™ Multicolor Broad Range Protein Ladder (Thermo; # 26623). Proteins were transferred to a polyvinylidene difluoride membrane by semi-dry transfer with a TransBlot Turbo (Bio-Rad). In Figs 1C and 7C, total protein, pre-and post-transfer, was monitored using the stain-free fluorophore as described (Posch et al., 2013; Ward, 2015a). Membranes were washed in TBST and blocked in TBST+5% milk (TBST-M; Nestle Carnation Instant Nonfat Dry Milk, 25.6-Ounce, Amazon) for one hour at room temperature. Blots were rocked in primary antibodies in TBST-M overnight at 4°C and then washed 4×5min with TBST. For primary antibodies conjugated with horseradish peroxidase (HRP) the blots were developed after the last TBST wash. Otherwise, blots were incubated with HRP-conjugated secondary antibodies in TBST-M for one hour at room temperature followed by 4×5min TBST washes and then developed as described below. The primary antibody used for Figures 1C,D, 2A, 5B, and 6B was horseradish peroxidase (HRP) conjugated anti-FLAG M2 (Sigma-Aldrich, A8592-5×1MG, Lot #SLCB9703) at a 1:2000 dilution. Precision Protein StrepTactin-HRP Conjugate (Bio-Rad, #1610381, Lot #64426657) was included with the primary antibody at a 1:10,000 dilution to visualize the protein size standard during blot imaging for Figs 1C,D. For the blot in Fig. 2A, 5B, 6B, we used mouse anti-alpha-Tubulin 12G10 (Developmental Studies Hybridoma Bank, the “-c” concentrated supernatant at 1:2000 (Fig 2A, S7) or 1:4000 (Fig. 5B, 6B). The secondary antibodies were Digital anti-mouse (Kindle Biosciences LLC, R1005) diluted 1:10,000 (Fig. 2A, S7) or 1:20,000 (Fig. S6B,C). Blots were incubated for 5 minutes with 1 ml of Supersignal West Femto Maximum Sensitivity Substrate (Thermo Fisher Scientific, 34095) and imaged using the ‘chemi high-resolution’ setting on a Bio-Rad ChemiDoc MP System.

For the blots in Fig. 6C, 20 μl was loaded on to NuPAGE 4 to 12% Protein Gels with 7 μl of 4X NuPAGE LDS sample buffer as described by the manufacturer (Invitrogen: NP0322BOX) and run for 2 h at 100 V, transferred to 0.45 micron nitrocellulose membrane (Bio-Rad) at 4 °C for 90 min using Bio-Rad WET transfer system. Following transfer, membranes were blocked in Intercept blocking buffer (LI-COR: 927-60001) for 1 h. Later, the membrane was incubated overnight in 1° Ab (ChromoTek: 32F6) at 1:5,000 dilution at 4°C on rocker. Membranes were washed 3X with TBST and incubated in 2° Ab (Sigma 12-349) at 1:100,000 dilution at RT on rocker for 1 h and washed 3X with TBST. Blots were visualized with Supersignal West Femto detection kit (Thermo Fisher) using a LI-COR Odyssey imager. Similarly, to test the loading control, 10 μl of F1 samples were loaded and following transfer to nitrocellulose membrane, probed with monoclonal mouse anti-actin clone C4 (Thermofisher scientific: ICN691001).

### Bioinformatics

Fig. 3A was generated in R (R Core Team). Phase and amplitude in Table S2 were converted to x and y coordinates for each gene by calculating x = amplitude * cos(phase) and y = amplitude * sin(phase). For Fig. 3D and S5, L3 NHR-23 ChIP-Seq data was downloaded from http://www.modencode.org, accession ID modEncode_3837. Wig files were converted to bigwig format and loaded into the Bioconductor package SeqPlots (Huber et al., 2015; Stempor and Ahringer, 2016). A bed file with the start and end coordinates of all genes’ CDS was generated (Wormbase WS220) and loaded into SeqPlots. SeqPlots aligned all genes by the start and end position and scaled the coding sequence of each gene to 2 kb. The average signal over all aligned genes was calculated in 25 bp windows from 1 kb upstream of the start site to 1kb downstream of the end site. Peak calls for NHR-23 in L3 (Fig 4C) were obtained from GEO (accession numbers: GSM1183659, GSM1183660, GSM1183661, GSM118366). Only peak calls validated by both replicates were considered. Peaks were assigned to the following genome features: gene body (WormBase WS220), 1kb upstream of the start site of coding genes, between 1kb and 3kb upstream of the gene start site and between 3kb and 5kb upstream of the gene start site. The same intervals were chosen downstream of the gene end sites. An NHR-23 peak was assigned to a feature if it overlapped with it by at least 100bp. The peak analysis data is in Table S5. The total number of peaks in all bins was tallied and is presented in Table S2 and in Fig. 3D. For Fig. 3D and 4E the phase for the oscillating genes in Table S1 was converted to time during the molting cycle by assuming a 9 hour larval stage which makes each hour=40°. We used the phasing from Meeuse et al., 2023 with lethargus starting at 45° and ecdysis ending at 135°. The phase in hours was then plotted along the x axis and the oscillating genes’ amplitude was plotted on the y axis. The gene annotation was based on the Concise Description and Automated Description attributes downloaded from Wormbase.

### RNAi

RNAi feeding plates were made by melting a solidified bottle of MYOB and adding Carbenicillin (25 ug/ml) and IPTG (8mM) once cooled to 55□. dsRNA-expressing *E. coli* bacteria were streaked on LB plates with Ampicillin (100 µg/ml) and grown overnight at 37□. A single colony was picked into 25 ml LB with 100 µg/ml Ampicillin and 12.5 µg/ml Tetracycline and shaken overnight at 37□ at 220 RPMs. The next day, the liquid culture was pelleted and resuspended in 1.25 ml LB with 100 µg/ml Ampicillin resulting in a 20x concentration from the original overnight liquid culture. 90 µl of the liquid culture was spread per small RNAi plate and allowed to dry. The plates were incubated at room temperature in the dark for 3 days. 10 adult worms from each strain were allowed to lay eggs on plates spread with *E. coli* bacteria containing either the control plasmid L4440 or the *nhr-23* RNAi knockdown plasmid for 1-2 hours. The adults were removed and the eggs allowed to develop at 20□ for 2 or 3 days depending on the experiment. For tissue-specific RNAi experiments we used RNAi defective *rde-1* mutant animals carrying *rde-1* transgenes to rescue RNAi in specific tissues (Qadota et al., 2007) The one exception was strain MGH171 which used an intestinal specific rescue of *sid-1* in an RNAi defective *sid-1* mutant (Melo and Ruvkun, 2012).

### Statistical Analysis

Statistical tests and numbers of animals analyzed are detailed in figure legends.

#### Competing interests

The authors declare no competing or financial interests.

## Acknowledgements

We thank Chris Hammell, David Matus, and Julian Ceron Madrigal for their critical reading of the manuscript. We thank David Fay, Miles Pufall, Helge Grosshans, and Ali Shariati for helpful discussions. We thank Jennifer Adams for generation of initial *rol-6* and *bli-1* mNeonGreen knock-ins. Figure S3A was created with Biorender.com.

## Funding

This work was funded by the National Institutes of Health (NIH) National Institute of General Medical Sciences (NIGMS) [R00GM107345 and R01GM138701] to J.D.W.

M.P. and A.D.C. were supported by NIGMS R35GM134970. Some strains were provided by the Caenorhabditis Genetics Center, which is funded by the NIH Office of Research Infrastructure Programs [P40 OD010440]. L.C.J. was funded by an NIGMS training grant [T32 GM133391]. The anti-alpha tubulin 12G10 monoclonal antibody developed by J. Frankel and E.M. Nelson of the University of Iowa was obtained from the Developmental Studies Hybridoma Bank, created by the NICHD of the NIH and maintained at The University of Iowa, Department of Biology, Iowa City, IA 52242.

---

**Fig. S1.**
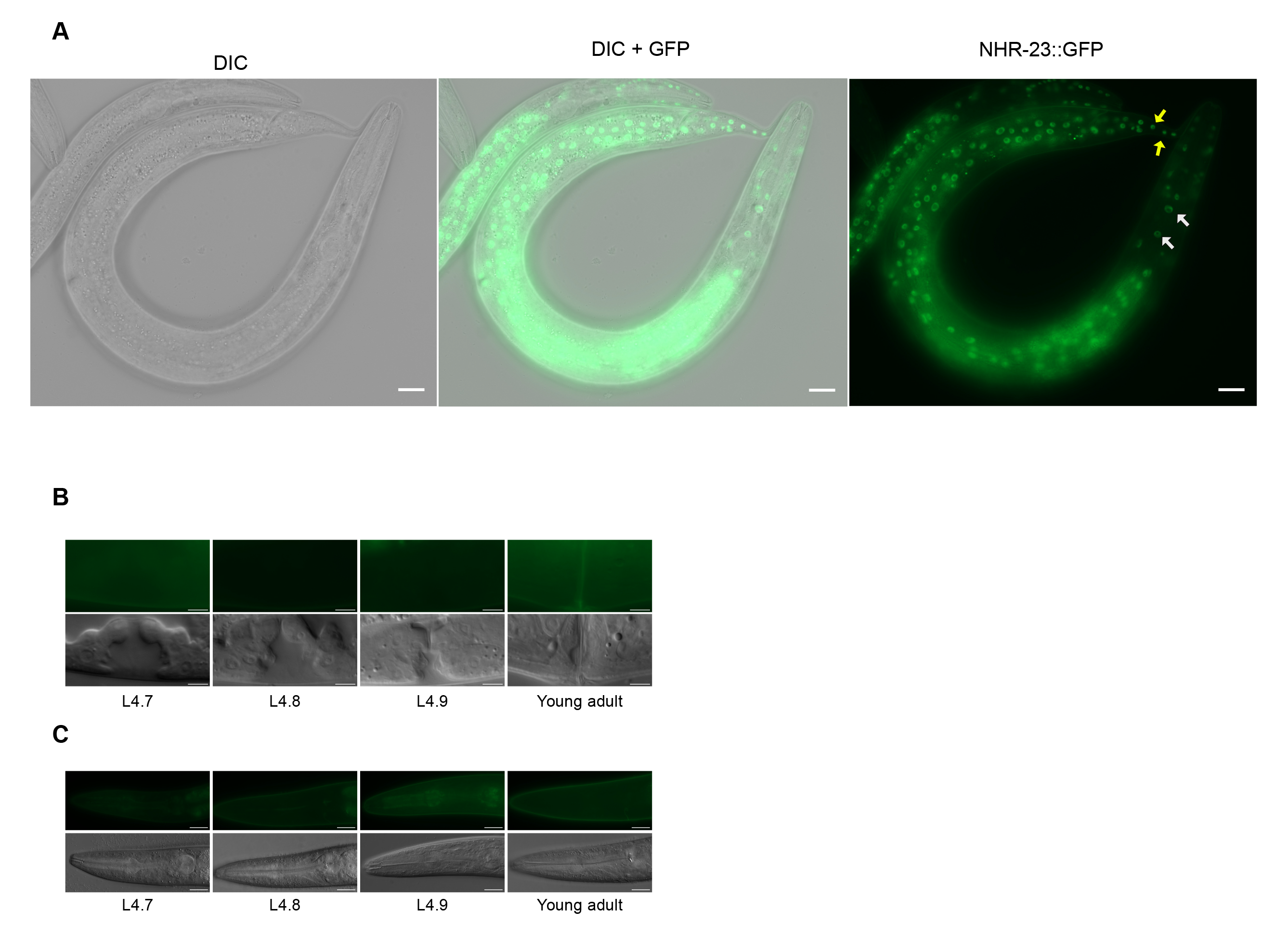
NHR-23::GFP expression. (A) Representative image of NHR-23 expression in *nhr-23::GFP::AID*::3xFLAG; TIR-1* L4 animals. NHR-23 expression is seen in the hypodermis and developing germline, as described in Ragle et al. 2020. White arrows point to seam cells expressing GFP and yellow arrows point to hyp cells expressing GFP. NHR-23 expression in the vulva (B) and head (C) of *nhr-23::GFP::AID*::3xFLAG* animals of the indicated stages. Images are representative of 20 animals examined over 4 independent experiments. Scale bars=5 µm (B) or 20 µm (C). Animals were staged based on vulval morphology (Mok et al., 2015).

**Fig. S2.**
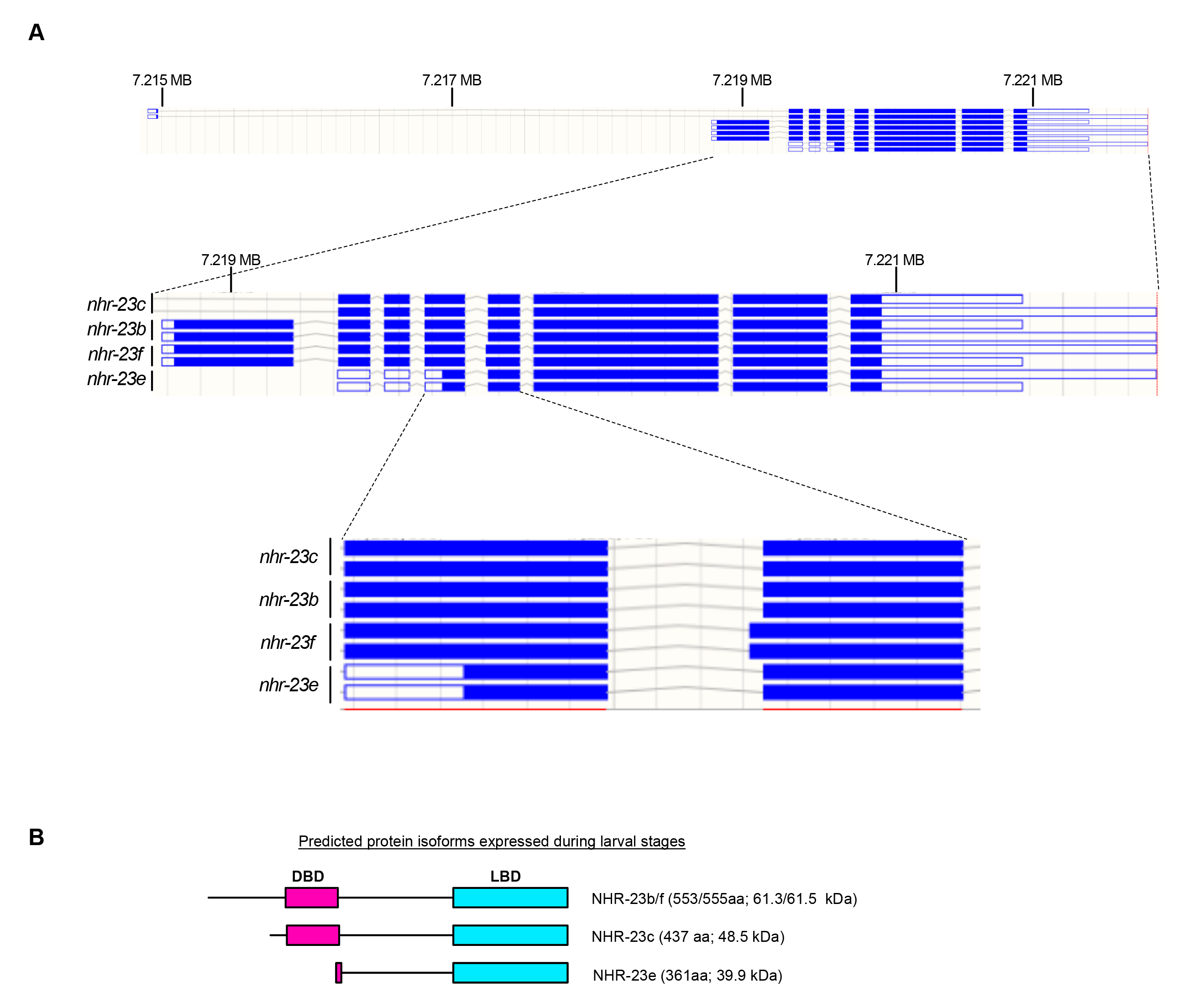
*nhr-23* isoforms detected by Nanopore direct mRNA sequencing. (A) Gene model diagrams of the *nhr-23* isoforms from the “observed isoforms” track on the ENSEMBL genome track. These are isoforms confirmed by direct mRNA sequencing (Roach et al., 2020). The top tracks are of the full *nhr-23* gene, including the *nhr-23c* isoforms, which contain a large first intron. The middle tracks are a zoomed in view starting at the *nhr-23b* transcription start site. The bottom tracks are further zoomed in to depict the alternative 3’ splice site usage that generates *nhr-23b* and *nhr-23f*. **(B)** Cartoons of predicted protein isoforms expressed during larval development based on Nanopore direct mRNA sequencing data (Roach et al., 2020). All of the predicted isoforms contain a Ligand Binding Domain (LBD) while only two of the three have a DNA Binding Domain (DBD). The predicted size of each isoform is listed to the right of the structure in amino acids (aa) and kDa. The AID*::3xFLAG fusion is predicted to add 9 kDa.

**Fig. S3.**
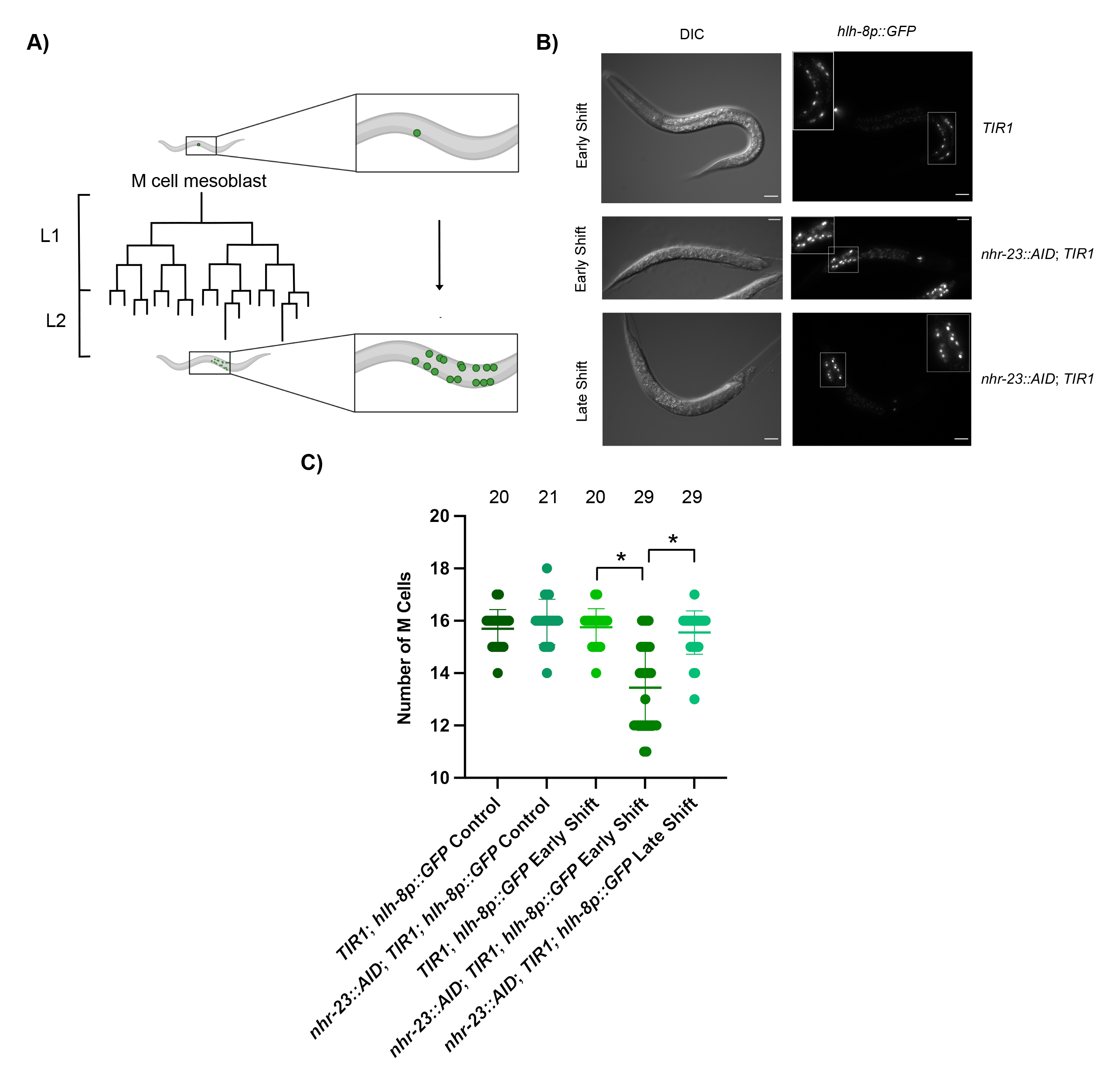
NHR-23 depletion causes developmental delay. (A) Depiction of the M cell mesoblast cell divisions in L1 larvae created with BioRender.com. Newly hatched animals are born with a single M cell, which undergoes the indicated set of divisions to produce 16 descendent cells at the L1/L2 molt. An *hlh-8p::GFP* reporter can be used to monitor M cell divisions in living animals. (B and C) Animals of the indicated genotype were synchronized and shifted to either auxin or control plates at either 3 (early shift) or 9 hours (late shift) post-release and imaged at 24 hours post-release. Two biological replicates were performed. (B) Representative images of animals of the indicated genotypes on auxin plates. (C) M cell quantification for animals of the indicated genotype shifted to control or auxin plates. Mean and standard deviation are labeled. The number of animals assayed for each genotype and condition is indicated at the top of the graph (* two-tailed T-test p<0.0001).

**Fig. S4.**
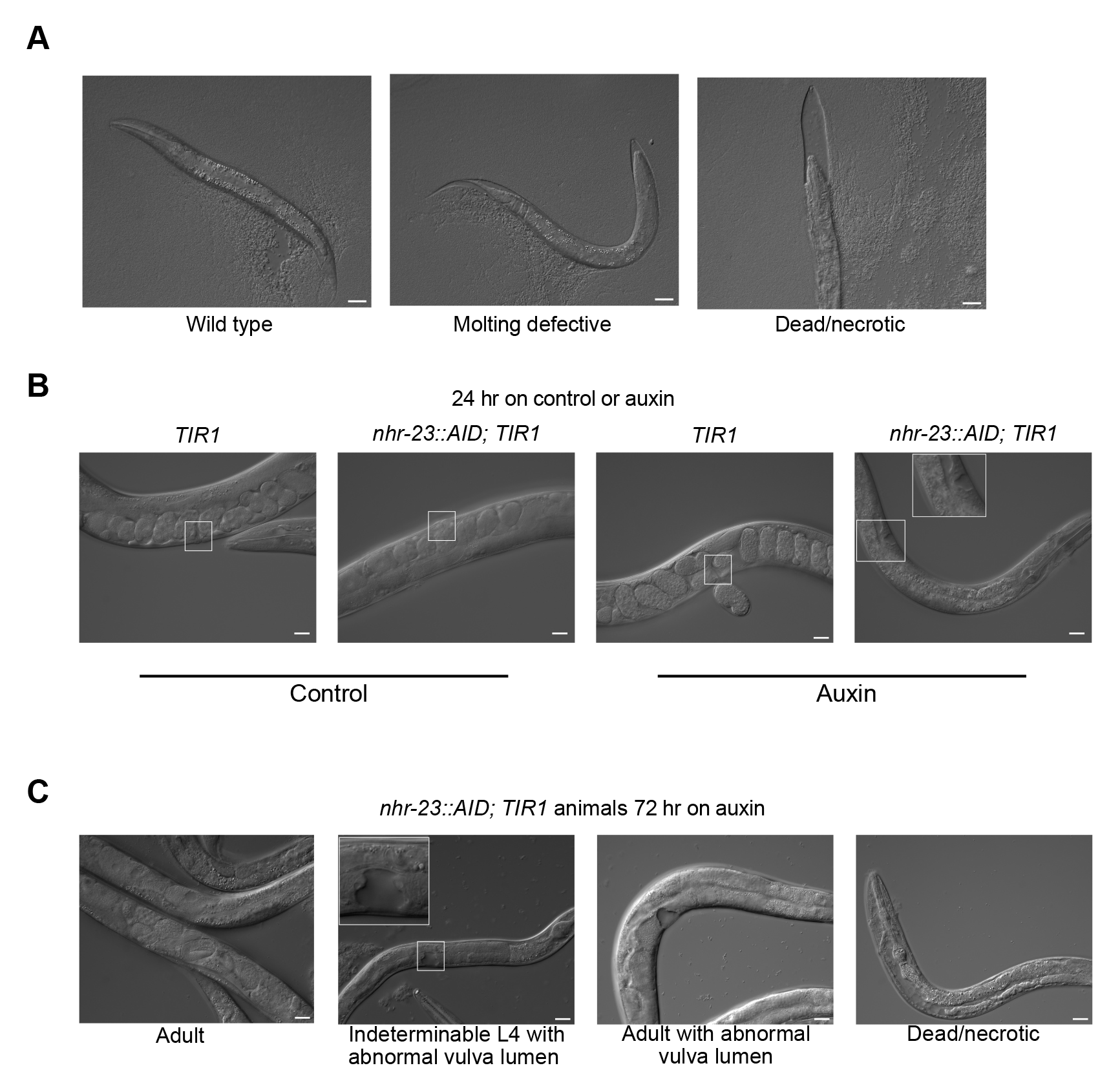
NHR-23 depletion causes developmental delay. (A) Representative images of synchronized *nhr-23::AID; TIR1* L1 larvae released on auxin and scored for molting defects and death/necrosis 72 hours post-release. (B) Representative images of animals of the indicated genotype that were synchronized and shifted onto auxin at 25 hours post-release and imaged 23 hours later (48 hours post-release). The position of the vulva is highlighted with a white box. (C) Representative images of *nhr-23::AID; TIR1* animals treated as in (B) and scored after 72 hours on auxin. Scale bars=20 µm. A zoomed inset is provided for the auxin-treated *nhr-23::AID, TIR1* images in B and C to highlight vulval morphology.

**Fig. S5.**
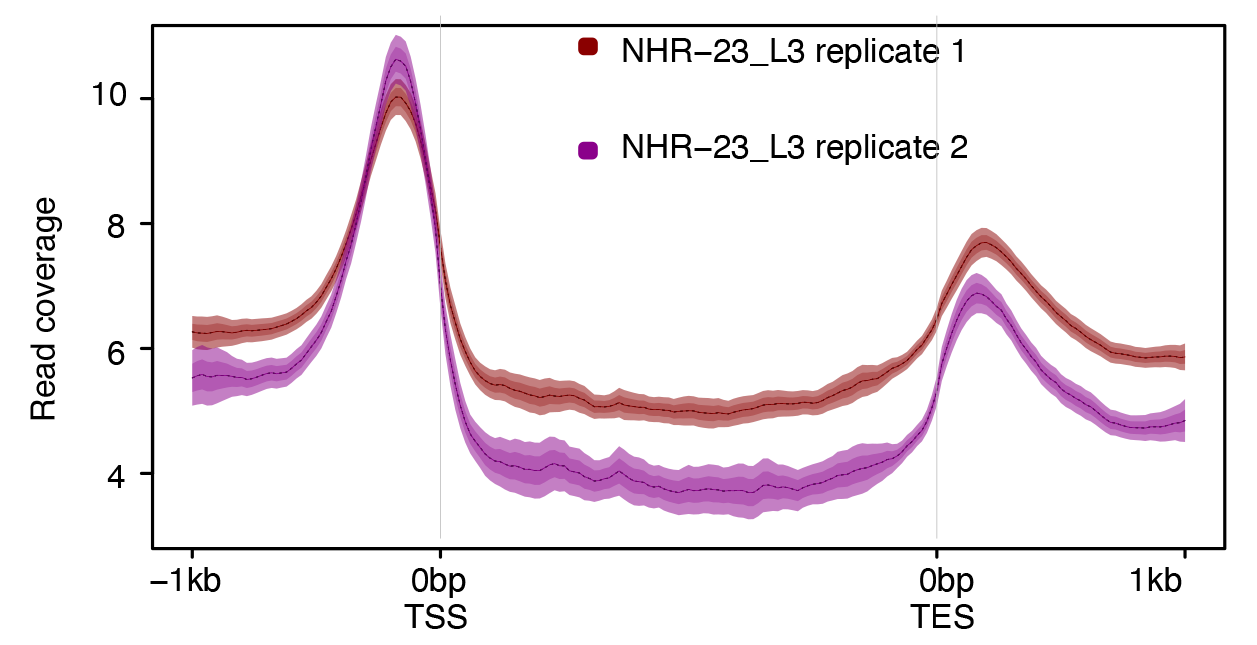
NHR-23 enrichment around genes. Average signal over all aligned genes from two NHR-23 ChIP-seq replicates (Gerstein et al., 2010). The y-axis indicates non-normalized read coverage. The mean signal is plotted with a line and the mean’s 95% confidence interval is indicated by the shaded area. TSS=transcription start site; TES=transcription end site.

**Fig. S6.**
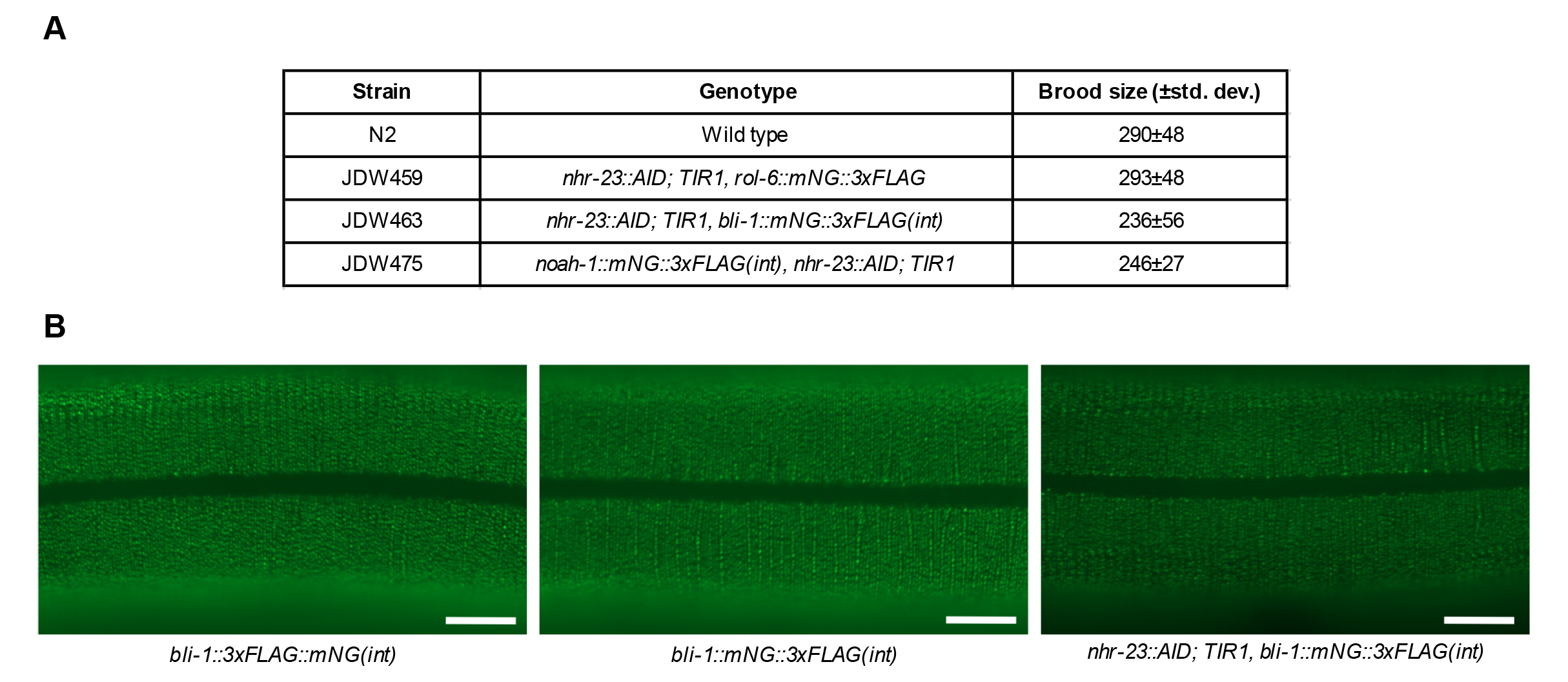
Validation of *bli-1, noah-1,* and *rol-6* knock-in strains. (A) Brood size analysis of animals of the indicated genotype. (B) Representative images from L4.9 stage CZ27591 *bli-1(ju1789[bli-1::3xFLAG::mNG(int)]) II*, JDW389 *bli-1(wrd84[bli-1::mNG::3xFLAG(int)]) II*, and JDW463 *nhr-23(kry61(nhr-23::AID*-TEV-3xFLAG)) I; ieSi57 [Peft-3::TIR1::mRuby::unc-54 3’UTR, cb-unc-119(+)], bli-1(wrd84[bli-1::mNG::3xFLAG(int)])* animals. Equal 500 msec exposures from the same area of animals were taken using a 100x objective. Knock-ins are all at the same location in *bli-1* (see Fig. 6A). Scale bars=10 µm. Images are representative of 40 animals examined in two independent experiments.

**Fig. S7.**
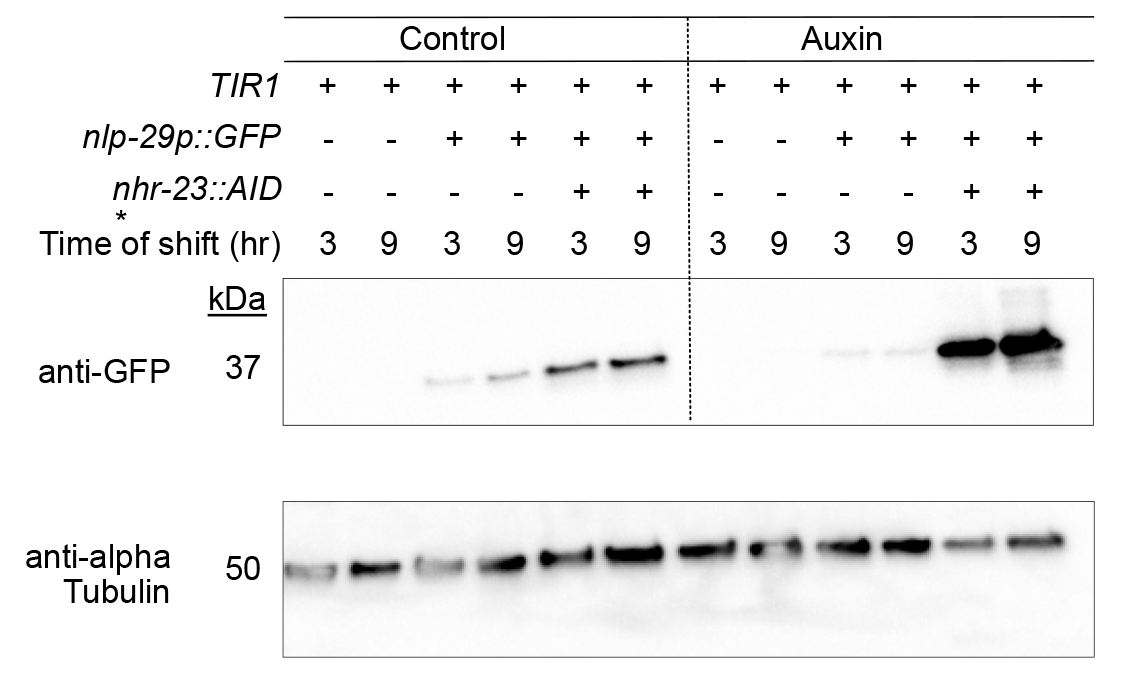
Depletion of NHR-23 leads to a elevated *nlp-29p::GFP* reporter activity. Anti-GFP immunoblot analysis of *nlp-29::*GFP expression. Marker size (in kDa) is provided. An anti-tubulin immunoblot is provided as a loading control. The blots are representative of two experimental replicates.

**Fig. S8.**
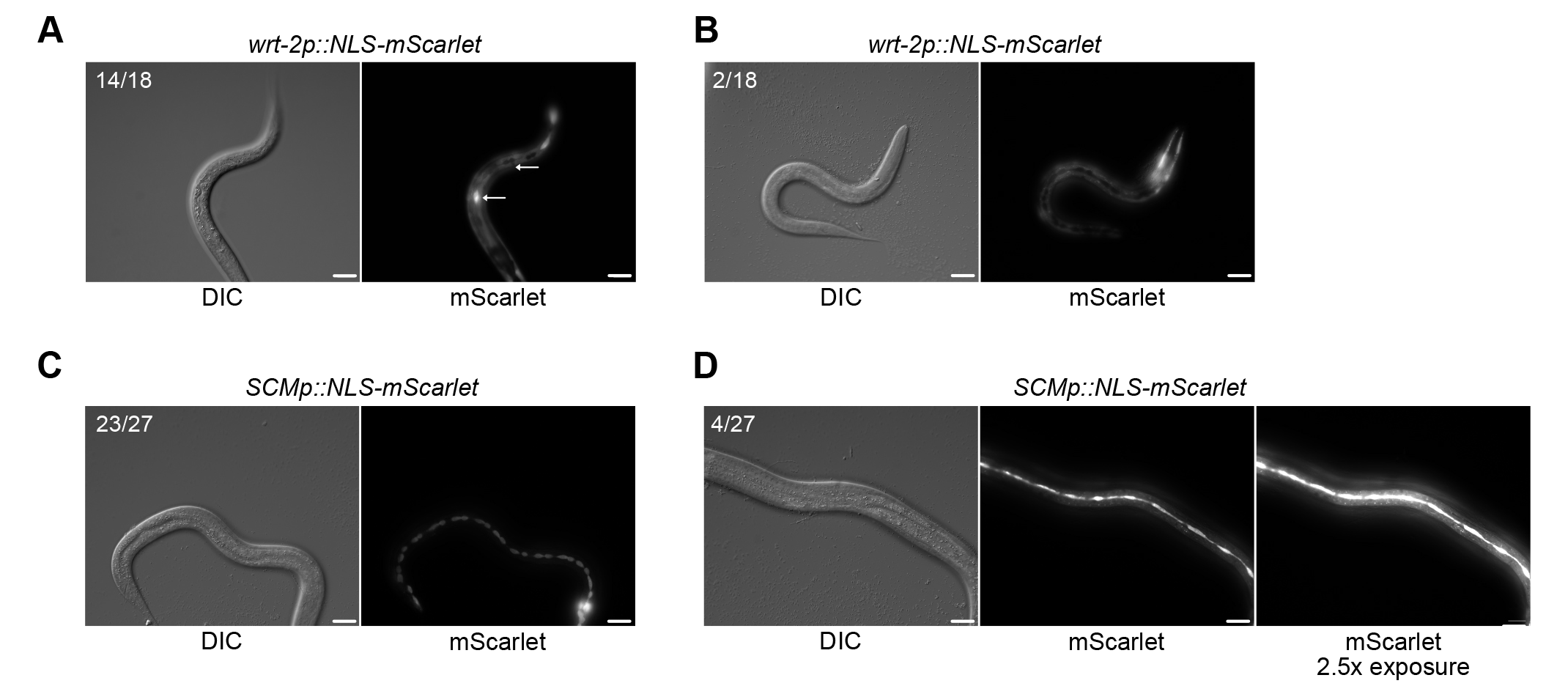
Expression patterns of *SCMp* and *wrt-2* promoter reporters. *wrt-2p::NLS::mScarlet* is expressed in the hypodermis and seam cells (A and B). In some animals seam cell expression is lacking. In A, arrows point to two seam cells, one with reporter expression and one lacking it. *SCMp::NLS::mScarlet* promoter reporters express robustly in the seam cells (C) with weak expression in the hypodermis (D) in a subset of animals. Two biological replicates were performed and the number of animals for which each image is representative is indicated.

**Fig. S9.**
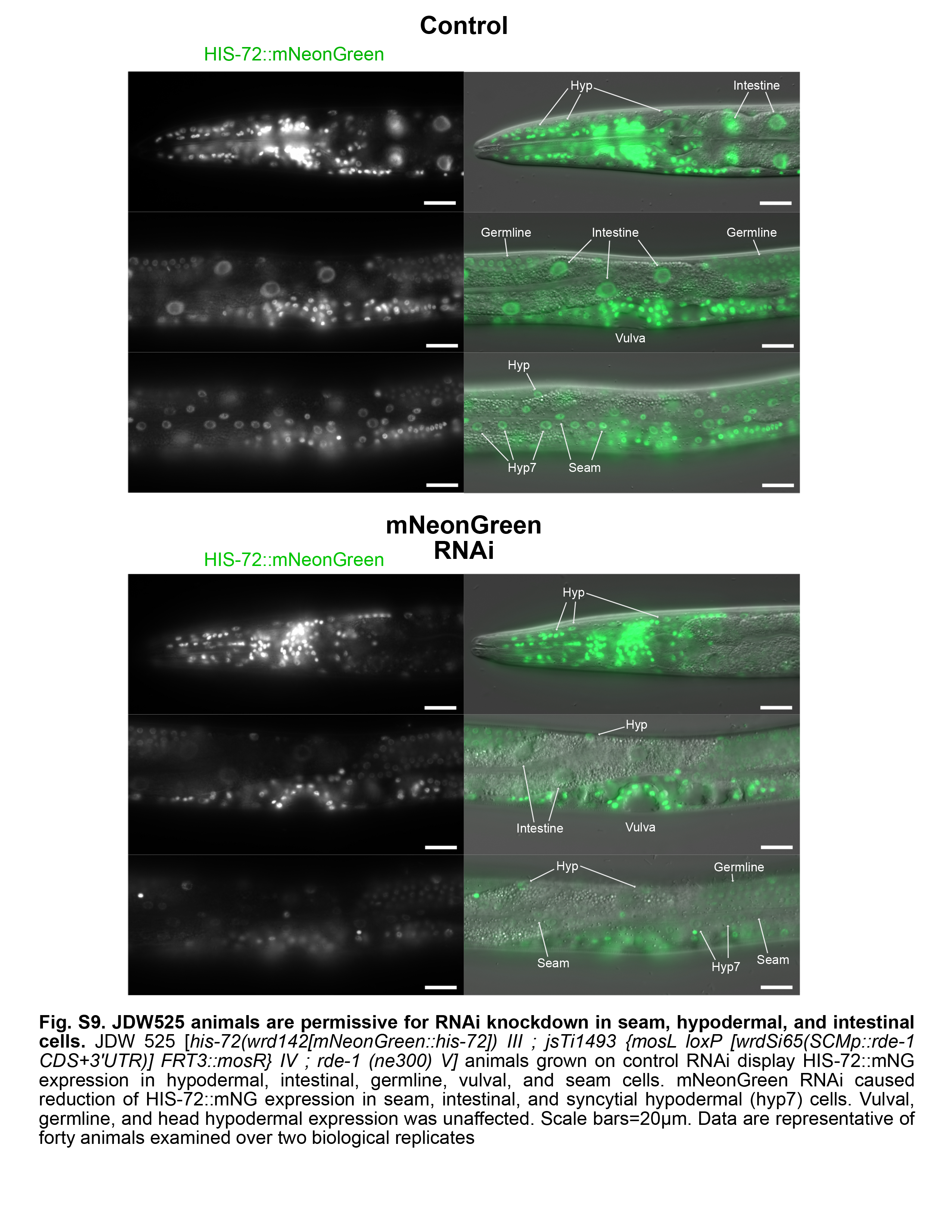
JDW525 animals are permissive for RNAi knockdown in seam, hypodermal, and intestinal cells. JDW 525 [his-72(wrd142[mNeonGreen::his-72]) III; jsTi1493 {mosL loxP [wrdSi65(SCMp::rde-1 CDS+3’UTR)] FRT3::mosR} IV; rde-1 (ne300) V] animals grown on control RNAi display HIS-72::mNG expression in hypodermal, intestinal, germline, vulval, and seam cells. mNeonGreen RNAi caused reduction of HIS-72::mNG expression in seam, intestinal, and syncytial hypodermal (hyp7) cells. Vulval, germline, and head hypodermal expression was unaffected. Scale bars=20µm. Data are representative of forty animals examined over two biological replicates

## Notes

### Competing Interest Statement

The authors have declared no competing interest.

### Summary of Updates

The text has been condensed, additional timecourse data and characterization of translational fusion strains has been provided.

